# Chromid-like secondary replicons as key sites of biosynthetic gene clusters in *Ktedonobacteria*

**DOI:** 10.64898/2026.02.02.703402

**Authors:** Shuhei Yabe, Yu Zheng, Shunji Takahashi, Chongyang Yang, Yui Nose, Shinichi Yamazaki, Nao Okuma, Mazytha Kinanti Rachmania, Fitria Ningsih, Wellyzar Sjamsuridzal, Mayuko Sato, Kiminori Toyooka, Yasunori Ichihashi

## Abstract

Soils harbour immense biosynthetic gene cluster (BGC) diversity that can mediate microbial interactions, yet this potential is still mapped unevenly across the tree of life. *Ktedonobacteria*—a class of actinomycete-like bacteria within phylum *Chloroflexota*—are widespread in terrestrial environments and repeatedly dominate pioneer communities in extremely oligotrophic volcanic bare-ground soils; however, their secondary metabolism and genome architecture remain poorly characterised. Here, we integrate targeted cultivation using volcanic soils from Mount Zao with genome-resolved metagenomics and public genomes to analyse 183 ktedonobacterial genomes. Using antiSMASH and BiG-SLiCE, we identified 1,546 BGCs comprising 1,162 non-redundant gene-cluster families (GCFs). In our dataset, nearly one quarter of genomes encode ≥10 distinct GCFs, and several family-level clades show mean GCF counts comparable to those in genus *Streptomyces*. Most ktedonobacterial BGCs are highly divergent from reference collections and exhibit unusually low intra-genomic redundancy, suggesting broad, underexplored chemotypes. Long-read assemblies from ten strains reveal recurrent 1.6–3.5 Mb chromid-like secondary replicons with chromosome-like composition but distinct maintenance signatures. These replicons are consistently enriched in BGCs and mobility-associated genes, with mobility loci concentrated near BGC boundaries. Collectively, our results expand the current knowledge of the phylogenetic landscape of soil biosynthetic diversity and highlight chromid-like secondary replicons as major genomic reservoirs for specialised metabolism in *Ktedonobacteria*.

## Introduction

Microbial secondary metabolism underpins diverse ecological interactions, including antagonism, resource acquisition, signalling, and stress tolerance [1]. Soils harbour large reservoirs of biosynthetic gene clusters (BGCs) encoding chemically diverse metabolites, many of which are exploited as antibiotics, agrochemicals, and other bioactive compounds [2, 3]. Yet, the breadth of lineages with high biosynthetic potential and the genomic architectures supporting this potential remain incompletely characterised [4]. Extreme terrestrial habitats provide an informative setting because intense, fluctuating stresses may favour specialised metabolites that mediate niche construction and competition [5–8].

Within these extreme habitats, *Ktedonobacteria* (phylum *Chloroflexota*) are often abundant in pioneer microbial communities, especially in volcanic bare-ground soils where they likely contribute to early primary succession on oligotrophic substrates [9–13]. These environments impose strong resource limitations and are characterised by fluctuating redox and climatic stresses, likely favouring flexible metabolic and defensive repertoires. *Ktedonobacteria* also exhibit a morphology akin to that of *Actinomycetes*, including filamentous growth, spores, and sporangia [14, 15], suggesting a potential coupling between complex life cycles and secondary metabolism.

Despite these features, the biosynthetic and ecological capacities of *Ktedonobacteria* remain underexplored compared with those of canonical discovery lineages such as *Streptomyces*. Individual strains have yielded unusual metabolites [15]. For example, *Thermosporothrix hazakensis* produces rare acyloins and indole–thiazole derivatives [16]. One of these compounds was previously optimised into ITE CONHCH₃, a synthetic aryl hydrocarbon receptor agonist effective in a murine colitis model [17]. However, such examples provide only a fragmentary view of the chemical potential of *Ktedonobacteria*, and little is known about how secondary metabolites contribute to their ecology in situ. Cultured representatives remain scarce, and isolate genomes have increased only modestly in recent years [18], whereas hundreds of metagenome-assembled genomes (MAGs) for *Ktedonobacteria* are now available. This shift expands phylogenetic coverage, but MAG fragmentation and incomplete recovery of secondary replicons can obscure genome architecture and secondary metabolism [19, 20].

One dimension of genome architecture relevant to secondary metabolism is the presence of large extrachromosomal elements (ECEs), which can concentrate secondary metabolism–related genes and facilitate the horizontal transfer and diversification of biosynthetic repertoires [21–23]. In *Streptomyces* and other bacterial genera, large ECEs (often classified as megaplasmids or chromids) serve as repositories of BGCs and underpin lineage-specific chemistry [23]. Chromids combine plasmid-like replication/maintenance with chromosome (CHR)-like features (e.g., similar GC content/codon usage and carriage of core genes) and have been proposed to facilitate niche adaptation [22, 24]. Several ECEs occur across multiple bacterial phyla [22, 25]; however, their prevalence and functional roles within *Ktedonobacteria* remain poorly defined. In a previous study, we noted ECE-like contigs in several *Dictyobacteraceae* strains, but their signatures and biosynthetic contributions remained unresolved [15]. In the present study, we combine targeted cultivation of volcanic bare-ground soil samples from Mount Zao (Japan) with genome-resolved metagenomics and public genome analysis to assess secondary metabolism and replicon architecture across *Ktedonobacteria*. Our findings showed that BGC diversity is widespread across this class, that chromid-like secondary replicons recur across multiple families and are enriched in BGCs and mobility-related genes, and that biosynthetic repertoires vary with lineage and habitat.

## Materials and Methods

### Soil sampling and TOC quantification

Samples of volcanic barren soil were collected on 29 September 2023 from the crater rim of Lake Okama, Mount Zao, Miyagi Prefecture, Japan (38.1321°N, 140.4468°E; 1,680 m above sea level; Supplementary Fig. S1A). Fifteen surface soil samples (0–10 cm) were collected using sterile tools and transported on ice to the laboratory within 48 h. Based on vegetation cover and total organic carbon (TOC) content, the samples were classified as non-vegetated soil (NVS; n = 10, 0.25%–0.41% TOC), poorly vegetated soil (PVS; n = 3, 0.46%–0.99% TOC), and richly vegetated soil (RVS; n = 2, 2.04% TOC). Subsamples were stored at –20°C for DNA extraction and at 4°C or –80°C for cultivation. TOC was quantified as described in Supplementary Methods.

### DNA extraction, sequencing, and assembly and MAG processing

Selected soil samples were subjected to amplicon (16S rRNA gene) sequencing and shotgun metagenomic sequencing, and reads were processed using MetaWrap pipelines [26]. Metagenomes were assembled and binned to recover MAGs, which were quality-filtered based on completeness and contamination thresholds and taxonomically assigned using GTDB-Tk [27]. Detailed protocols, software versions, and parameters are provided in Supplementary Methods.

### Isolation, genome sequencing, and scanning electron microscopic (SEM) analysis of *Ktedonobacteria*

The isolation, identification, complete genome sequencing, and SEM analysis of *Ktedonobacteria* are described in Supplementary Methods. Briefly, isolates were obtained on acidified 1/10-strength R2A–gellan and identified by near full-length 16S rRNA gene sequencing and the EZBioCloud bioinformatics platform [28]. Two strains were fully assembled from long reads (Flye) [29] and polished/circularised using the Pilon [30] and Circlator [31] tools.

### Phylogenomics of *Ktedonobacteria*

The procedures for phylogenomic analysis are described in detail in Supplementary Methods. Briefly, a dataset comprising 184 ktedonobacterial genomes was compiled by combining public genomes from the National Center for Biotechnology Information with 21 MAGs derived from Mount Zao samples and two new isolates (Z7_2 and Z4_9). Public MAGs were dereplicated at 95% average nucleotide identity (ANI) using dRep (fastANI) [32]. Phylogeny was inferred from GTDB-Tk bac120 marker genes using maximum-likelihood in IQ-TREE after standard alignment and trimming (Supplementary Methods). Family-level clades were delineated with TreeCluster (“max_clade”) using a distance threshold of 0.45 [33], and clade assignments were subsequently integrated with BGC profiles and replicon architecture in downstream comparative analyses.

### BGC prediction, clustering, diversity, and novelty analysis

BGC prediction and comparative analyses are described in detail in Supplementary Methods. Briefly, the analysis included a total of 183 ktedonobacterial genomes (excluding Merged_NVS_bin65), and BGCs were predicted using antiSMASH v7.1.0 [34], restricting the input to scaffolds ≥ 5 kb to limit fragmentation bias. BGCs were clustered into gene cluster families (GCFs) using BiG-SLiCE v2.0 at a cosine of 0.4 [35], and GCF counts were mapped onto the phylogeny (iTOL) [36]. Following the metric recommended by Gavriilidou et al. [37], lineages were defined as BGC-rich when genomes harboured ≥10 distinct GCFs, and clades were defined as highly BGC-rich when the mean per-genome GCF count across member genomes was ≥15. GCFs were assigned to major biosynthetic classes based on antiSMASH/BiG-SLiCE annotations. Comparisons across lineages and genome types were tested using Wilcoxon rank-sum tests with Benjamini–Hochberg correction. Novelty was evaluated using (i) BiG-SLiCE v1.1.1 against the BiG-FAM dataset [38], (ii) BiG-SCAPE against the MiBIG v4.0 database [39], and (iii) antiSMASH KnownClusterBlast [34].

### Identification and comparative analysis of ECE-like contigs

Candidate ECE-like contigs were identified from high-quality long-read assemblies (≤10 contigs) using operational criteria (Supplementary Methods). Briefly, contigs ≥1 Mb long were screened as putative large replicons based on the reported size distributions of chromids versus megaplasmids [22]. To distinguish large ECEs from fragmented CHRs, we compared observed CheckM completeness (domain Bacteria) to the expected value proportional to contig length (contig length / total genome size) and designated contigs as ECE-like when the normalised completeness was ≤0.2 (Supplementary Methods).

Replication-related features (e.g. strand asymmetry, predicted origins, and replication/partition genes) were profiled, and ECE-like contigs were compared with their cognate CHR-like contigs using genome similarity, orthology-based comparisons, and functional annotation (Supplementary Methods). To test whether mobility-associated genes were depleted within BGCs and/or enriched near BGC boundaries, we quantified overlap and proximity between antiSMASH-predicted BGC intervals and mobility-gene intervals and evaluated enrichment using length-corrected tests complemented by within-replicon permutations (Supplementary Methods). Contig-scale synteny and feature tracks were visualised using Circos [40].

## Results

### Distribution and recovery of *Ktedonobacteria* from Zao Volcano soils

The Okama crater rim at Mount Zao is characterised by oligotrophic volcanic bare-ground soils interspersed with localised vegetated patches, providing a natural gradient of early-stage soil development (Supplementary Fig. S1A). To investigate the distribution of *Ktedonobacteria*, we conducted 16S rRNA gene (V4 region) amplicon sequencing on 15 soil samples categorised by vegetation cover into NVS, PVS, and RVS (Supplementary Data S1). *Ktedonobacteria* represented one of the dominant bacterial classes across samples (Supplementary Fig. S1B). Notably, their relative abundance tended to decrease with increasing vegetation cover, with ranges of 3.0%–17.6%, 0.60%–12.22%, and 0.03%–4.94% in NVS, PVS, and RVS, respectively. This pattern was consistent with the preference for bare-ground soils observed in this dataset, although the small sizes of PVS and RVS samples preclude statistical inference (Supplementary Fig. S1C).

Metagenomic sequencing of five NVS and three PVS samples yielded a total of 166 MAGs (≥50% completeness, <10% contamination) spanning 12 bacterial phyla [41]. dRep clustering (≥95% ANI) identified 90 species-level clusters, including 21 ktedonobacterial MAGs grouped into nine species-level clusters (Supplementary Data S2, S3) [42].

We also attempted cultivation of *Ktedonobacteria* from the same samples. The colonies were screened on modified R2A medium under oligotrophic conditions, using pH adjustment and antibiotics to suppress fast-growing heterotrophs. Colonies with substrate-rooted mycelia were selected and identified via 16S rRNA gene sequencing. Among 56 colonies examined, four were assigned to *Ktedonobacteria*. After three rounds of purification, four axenic strains representing two species were obtained, designated as strains Z7_2 (JCM 37047; DSM 118755) and Z4_9 (JCM 37048; DSM 118756), respectively (Supplementary Data S4).

### Phylogenetic structure of *Ktedonobacteria* based on MAGs and cultured genomes

A maximum-likelihood phylogeny was constructed using the GTDB bac120 marker set (35,833 aligned amino acid positions) (Fig. 1A). The dataset included 21 MAGs from Mount Zao samples, 21 genomes of cultivated strains, the newly isolated strains Z7_2 and Z4_9, and 140 representative MAGs selected from 296 public ktedonobacterial MAGs (as of May 2025). Public MAGs were dereplicated at ≥95% ANI to retain one representative per species-level cluster (Supplementary Data S5).

**Fig. 1.**
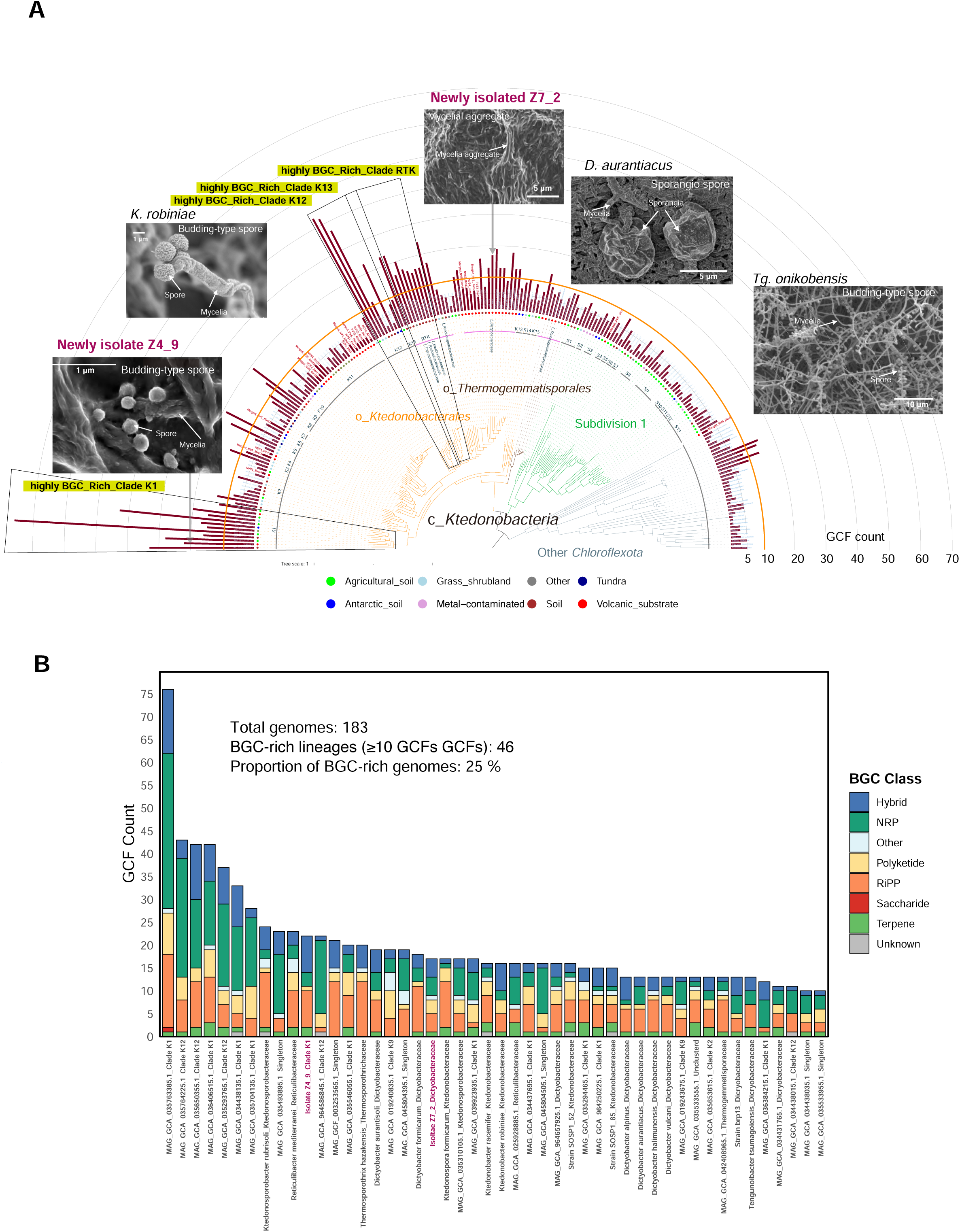
Phylogenetic structure, habitat distribution, and biosynthetic capacity of *Ktedonobacteria*. **(A)** Maximum-likelihood phylogeny inferred from the GTDB bac120 marker set (35,833 aligned amino acid positions) for 183 genomes (metagenome-assembled genomes, previously cultivated representatives, and the newly isolated strains Z7_2 and Z4_9 from Mount Zao samples). Order-level clades are colour-coded. Radial bars denote the number of GCFs per genome (antiSMASH v7.1.0 calls clustered with BiG-SLiCE 2.0 at cosine 0.40). Point symbols indicate habitat categories as shown in the legend. Family-level clades within *Ktedonobacterales* (e.g. K1, K12, K13, and RTK) were delineated using TreeCluster (v1.0.4, max_clade mode, threshold 0.4). BGC-rich lineages were defined as genomes with ≥10 GCFs, and highly BGC-rich clades as family-level groups with ≥15 GCFs per genome. Representative scanning electron micrographs illustrate that actinomycete-like filamentous morphology is broadly conserved across distinct clades (scale bars as indicated). **(B)** Distribution of biosynthetic gene cluster classes across the 183 ktedonobacterial genomes examined. Bars show per-genome GCF counts, stacked by antiSMASH class and ordered by total GCFs. The inset summarises the total number of genomes, the number of BGC-rich lineages (≥10 GCFs per genome), and their proportion. GCF, gene-cluster family; BGC, biosynthetic gene cluster.

The obtained phylogeny resolved three major lineages—*Ktedonobacterales*, *Thermogemmatisporales*, and an uncultivated “Subdivision 1” (Fig. 1A). TreeCluster (v1.0.4; max_clade; threshold = 0.4) was applied to define 37 operational phylogenetic clusters for downstream analyses (Supplementary Data S5), denoted as Clade K (*Ktedonobacterales*) and Clade S (Subdivision 1) in Fig. 1A. Under this parameterisation, *Ktedonobacteraceae*, *Reticulibacteraceae*, and *Thermosporotrichaceae* were merged as Clade RTK (formal family names retained in taxonomic contexts). Habitat information, antiSMASH-predicted BGC counts, and representative SEM images were mapped onto the tree (Fig. 1A). Twenty out of the 21 MAGs from Mount Zao samples corresponded to defined ktenodonobacterial lineages, including Clades K2, K4, and K11, two novel singleton lineages, and the known *Dictyobacteraceae* family. One MAG (Merged_NVS_bin65) fell outside the core *Ktedonobacteria* clade (Fig. 1A), forming a distinct branch basal to other *Chloroflexota* taxa, and it was therefore excluded from downstream BGC analyses.

The newly isolated strain Z7_2 was grouped with *Dictyobacter aurantisoli* Uno17 but shared only 95.43% 16S rRNA gene identity (vs. the 98.65% species threshold[43]), supporting its recognition as a previously undescribed species within *Dictyobacteraceae*. In contrast, Z4_9 fell within the uncultivated Clade K and showed 90.25% 16S rRNA identity to *D. aurantisoli*, consistent with a deeply branching lineage at approximately family-level divergence (Supplementary Data S4).

### Biosynthetic potential of *Ktedonobacteria* compared with Streptomyces

The secondary metabolic potential of 183 ktedonobacterial genomes (22 cultured, 161 MAGs) was assessed using antiSMASH v7.1.0, resulting in the identification of 1,546 BGCs, which were dereplicated using BiG-SLiCE 2.0 (cosine threshold = 0.40) into 1,162 non-redundant GCFs (Supplementary Data S6). Using the same workflow for other *Chloroflexota* classes, *Streptomyces*, and representative *Actinomycetota*, we identified 46 species-level lineages (25%) as “BGC-rich” (≥10 GCFs per genome) (Fig. 1B). Within *Ktedonobacterales*, four TreeCluster clades (K1, K12, K13, and RTK) were highly BGC-rich (≥15 GCFs per genome), and the newly isolated strain Z4_9 was identified as belonging to the BGC-rich Clade K1.

Across genomes, *Ktedonobacteria* averaged 8.4 GCFs per genome (n = 183), nearly twice the number observed for other *Chloroflexota* classes (4.4; n = 38; p = 0.006) (Fig. 2A). Cultured *Ktedonobacteria* were particularly BGC-rich (14.7; n = 22), exceeding ktedonobacterial MAGs (7.5; n = 161; p = 3.2 × 10⁻⁷) (Supplementary Fig. S2) and actinomycetotal genomes (12.0; n = 1,020; p = 0.011). Notably, the most BGC-rich ktedonobacterial genomes exceeded the mean GCF count of *Streptomyces* (Fig. 2A). Within the *Ktedonobacteria* class, *Ktedonobacterales* and *Thermogemmatisporales* showed higher GCF counts than Subdivision 1 (Supplementary Data S7).

**Fig. 2.**
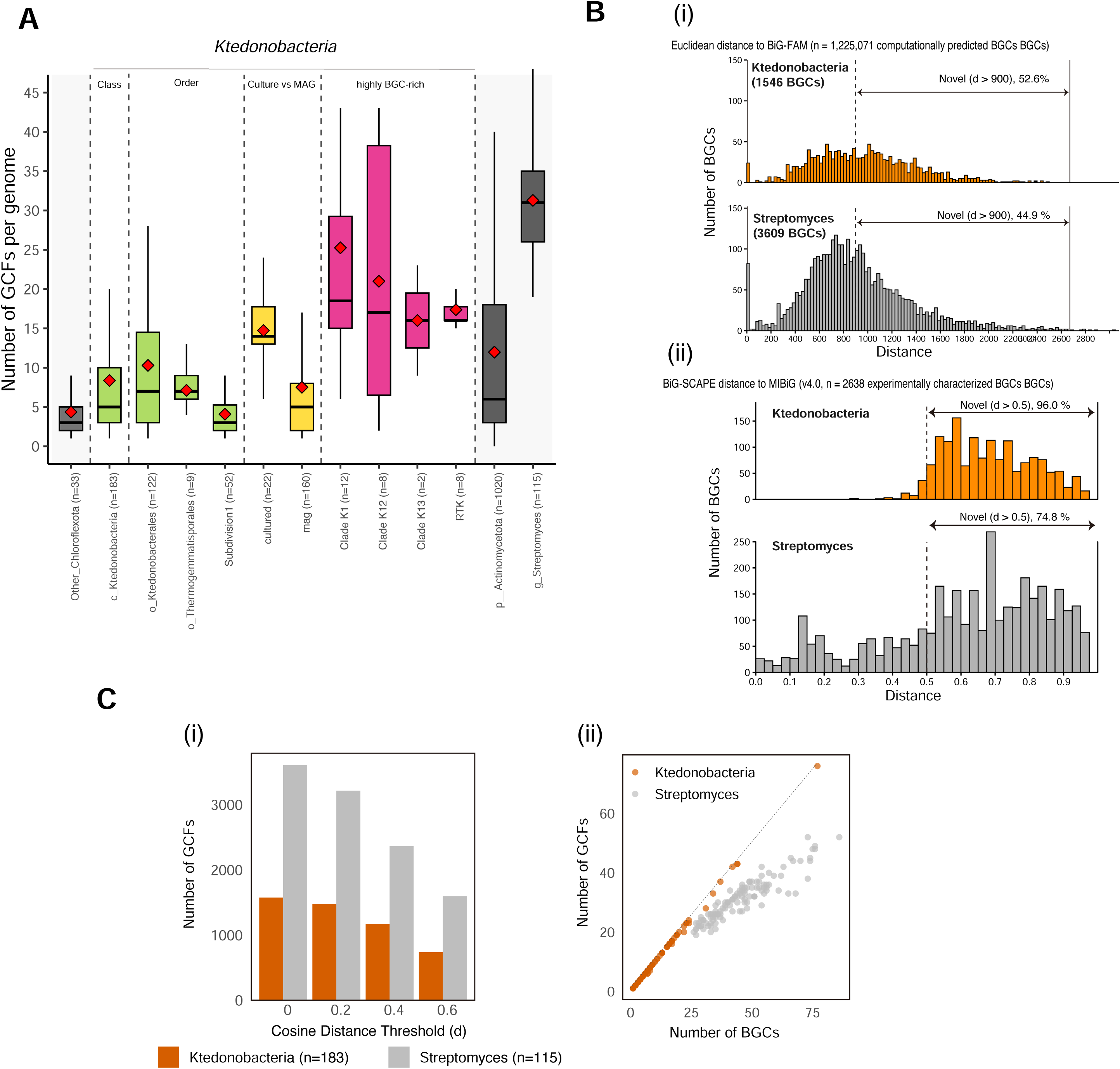
Comparative analysis of BGC abundance, distance-based novelty, and redundancy in *Ktedonobacteria* versus *Streptomyces*. **(A)** Number of GCFs per genome across groups. Box-and-whisker plots summarise per-genome GCF counts for orders within *Ktedonobacteria* (including cultured strains and MAGs), other classes of *Chloroflexota*, genus *Streptomyces*, and broader *Actinomycetota*. Boxes represent the IQR with the median line; whiskers extend to 1.5x IQR. Red diamonds indicate the mean value for each group. GCFs were obtained by running antiSMASH v7.1.0 and clustering the resulting BGCs with BiG-SLiCE 2.0 at a cosine of 0.40. Statistical comparisons are provided in Supplementary Data S7. **(B)** Distance-based novelty of BGCs. (i) Distribution of Euclidean distances to the BiG-FAM database dataset for the BGCs of *Ktedonobacteria* and *Streptomyces* (1,225,071 BGCs); larger distances indicate lower similarity to known families. (ii) Distribution of BiG-SCAPE distances (d) from each BGC to its closest MIBiG BGC; a higher d value indicates lower similarity. Proportions above the indicated thresholds are annotated in the panels. **(C)** Redundancy and threshold sensitivity. (i) Sensitivity of GCF counts to the BiG-SLiCE cosine threshold (results summarised across a range of thresholds). (ii) Relationship between the numbers of BGCs and GCFs per genome for *Ktedonobacteria* and *Streptomyces*, illustrating that ktedonobacterial genomes tend towards a near one-to-one correspondence (low redundancy), whereas *Streptomyces* genomes more often harbour multiple BGCs within the same GCF. GCF, gene-cluster family; BGC, biosynthetic gene cluster; IQR, interquartile range.

Novelty analyses indicated that ktedonobacterial BGCs diverged substantially from current references (Fig. 2B), with 52.6% showing low similarity (Euclidean distance > 900) against the BiG-FAM dataset, compared with 44.9% of BGCs showing low similarity in *Streptomyces* (Fig. 2B(i)). When using BiG-SCAPE against the MIBiG database, 96.0% of ktedonobacterial BGCs exceeded d = 0.5 (vs. 74.8% for *Streptomyces*) (Fig. 2B(ii)). Consistently, KnownClusterBlast reported <20% similarity for >80% of ktedonobacterial BGCs, with only 5% (80/1,546) showing ≥80% similarity to known clusters (Supplementary Data S8).

Varying the BiG-SLiCE cosine threshold (0.0–0.6) affected *Ktedonobacteria* and *Streptomyces* in similar ways (Fig. 2C(i)). However, the former showed a ratio of approximately 1:1 between BGC and GCF counts per genome, consistent with low intra-genomic redundancy, whereas the latter more often carried multiple BGCs within the same GCF (Fig. 2C(ii)).

Collectively, these results show that some ktedonobacterial clades have biosynthetic capacities comparable to or exceeding those of *Streptomyces* and that many BGCs diverge substantially from current references. Because database-based comparisons are limited by incomplete and biased coverage, chemical characterisation will be required to confirm novelty and function.

### Lineage- and habitat-associated structuring of biosynthetic repertoires

Uniform manifold approximation and projection of subclass-level BGC profiles separated genomes primarily by phylogenetic lineage and, to a lesser extent, by habitat (Supplementary Fig. S3; Supplementary Data S9). Habitat labels, which were coarse categories derived from BioSample metadata, were interpreted as broad associations rather than detailed environmental classifications. PERMANOVA confirmed significant clustering by lineage (pseudo-F = 11.2, R² = 0.71, p = 0.001) and by habitat (pseudo-F = 2.42, R² = 0.09, p = 0.011), indicating that evolutionary history explains most variance, with habitat contributing a smaller but detectable fraction. Lineage-level comparisons revealed distinct biosynthetic signatures (Fig. 3A).

**Fig. 3.**
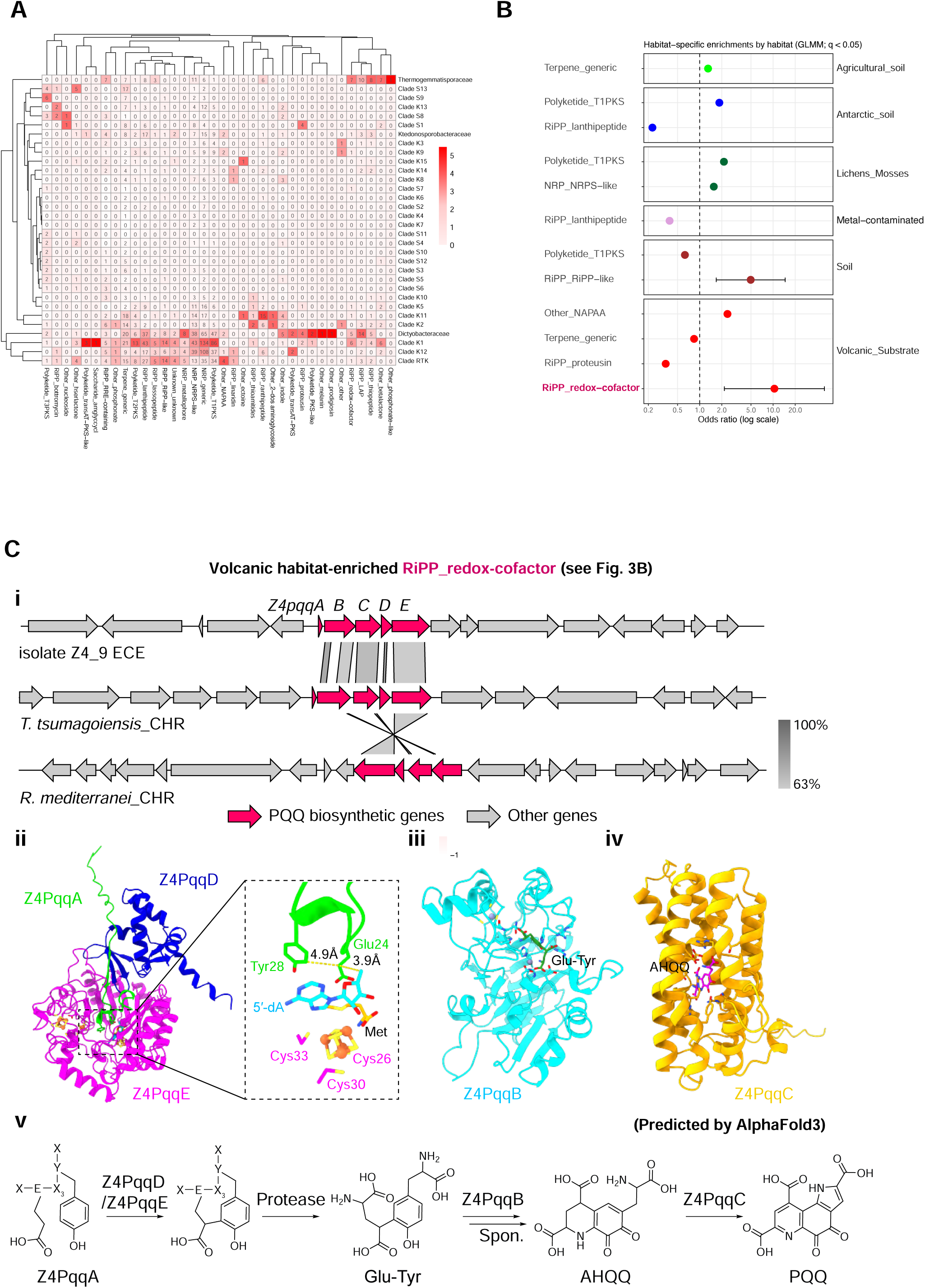
Lineage- and habitat-specific structuring of ktedonobacterial biosynthetic repertoires and a volcanic habitat-enriched RiPP_redox-cofactor pathway. **(A)** Heatmap showing the distribution of biosynthetic categories across taxonomic groups. Each row represents a taxonomic group and each column a biosynthetic category assigned at the GCF level (shown on the x-axis as “Class_Subclass”; e.g. RiPP_bottomycin). Genomes assigned to “Singleton” were excluded. To emphasise category-specific enrichment across taxa, values were Z-score-normalised per column, and original (untransformed) counts are overlaid as numbers in each cell. GCFs were classified using BiG-SLiCE, and biosynthetic categories were assigned based on antiSMASH/BiG-SLiCE annotations as described in Methods. For hybrid-type GCFs, all associated biosynthetic categories were counted independently. **(B)** Habitat-specific enrichment in biosynthetic categories identified by GLMM (Benjamini–Hochberg correction within habitat, q < 0.05). Odds ratios are shown with 95% confidence intervals. The category RiPP_redox-cofactor enriched in volcanic habitats is highlighted in red. Statistical details are provided in Supplementary Data S10. **(C)** Volcanic habitat–enriched RiPP_redox-cofactor locus and PQQ-like locus inference. (i) Comparative analysis of RiPP_redox-cofactor clusters annotated by antiSMASH from the ECE of strain Z4_9, CHR of *Thermogemmatispora tsumagoiensis*, and CHR of *Rhodococcus mediterranei*. Genes involved in PQQ biosynthesis (pqqA–E) are shown in red and other genes in grey; sequence identity is indicated by grey shading. (ii) AlphaFold 3-predicted complex model of Z4PqqA, Z4PqqD, and Z4PqqE with the SAM-derived products 5L-dA) and Met. The Glu24 and Tyr28 residues of the EXXXY motif in Z4PqqA are shown as sticks, and distances are indicated by yellow dashed lines. (iii) and (iv) AlphaFold 3-predicted complex models of Z4PqqB bound to the Glu–Tyr substrate (panel iii) and Z4PqqC bound to the AHQQ intermediate (panel iv). Key residues surrounding the ligands are shown as sticks. (v) Putative functional model for Z4PqqA–E in strain Z4_9 within a PQQ-like (peptide-derived redox-cofactor) locus, inferred from gene content and structure predictions. The proposed maturation scheme is based on previously established models [44]. mET, methionine; 5L-dA, 5L-deoxyadenosine; GLMM, generalised linear mixed model; GCF, gene-cluster family; BGC, biosynthetic gene cluster.

*Thermogemmatisporaceae* were enriched in RiPP subclasses (including thiopeptides and RRE-containing peptides) and betalactones, whereas Clade K11 exhibited a predominance of lantipeptides and terpenes and lacked polyketide synthase (PKS)-related clusters. *Dictyobacteraceae* uniquely encoded traits such as melanin/prodigiosin biosynthesis, whereas BGC-rich clades (K1, K12, and RTK) were enriched in PKS and non-ribosomal peptide synthetase clusters.

To account for the phylogenetic structure, we fitted GLMMs with lineage as a random effect (Fig. 3B; Supplementary Data S10) and detected several habitat-associated shifts in BGC subclasses. Soil-derived genomes, Antarctic/lichen-associated genomes, and volcanic substrates were enriched in RiPP_RiPP-like clusters (OR = 4.96, q = 0.044), Polyketide_T1PKS clusters (OR = 1.85–2.14), and RiPP_redox-cofactor clusters (OR = 10.4, q = 0.031), respectively. Given the coarse habitat labels and uneven sampling, these results were interpreted as indicative trends rather than strict habitat-specific signatures.

Consistent with this cautious interpretation, RiPP_redox-cofactor clusters occurred in the genomes from Mount Zao samples (including Z4_9) and in additional genomes from volcanic and non-volcanic habitats (Supplementary Data S9). These clusters showed low similarity to known references in KnownClusterBlast (13%–20% to lankacidin C) and contained conserved pqqA–E-like genes consistent with PQQ-like loci (Supplementary Data S8).

To further characterise RiPP_redox-cofactor clusters (Fig. 3B), we examined Z4_9 together with the type strains *Tengunoibacter tsumagoiensis* and *Reticulibacter mediterranei*. All three genomes were shown to harbour highly syntenic loci (Fig. 3C(i)) and, in Z4_9, the locus encodes pqqA–E in the canonical order. AlphaFold3 predictions were consistent with a canonical PQQ biosynthetic pathway, including a RiPP chaperone-like PqqD, a radical SAM-like PqqE, and PqqB/PqqC folds compatible with downstream oxidative steps (Fig. 3C(ii–v)). No pqqF homologue was identified within these loci, suggesting that the proteolysis required for maturation may be provided in trans [44]. Accordingly, a genome-encoded protease that complements the function of PqqF might be involved in PQQ production.

### Large chromid-like ECEs in *Ktedonobacteria*

To link BGCs with large ECEs, 20 ktedonobacterial genomes (18 type strains plus Z4_9 and Z7_2) were scanned for large contigs distinct from the primary CHR. For the initial screening, ECE-like contigs were defined operationally as ≥1 Mbp sequences whose CheckM-estimated completeness was ≥5-fold lower than expected based on their size relative to the total genome, which restricted the analysis to high-contiguity long-read assemblies (≤10 contigs) (Table 1; Supplementary Data S11; cf. chromid and megaplasmid size distributions in diCenzo & Finan [22]). Using this definition, ten genomes (eight type strains and both new isolates) with at least one such replicon were identified; the remainder had a single CHR, lacked ≥1 Mbp contigs, or were too fragmented to be evaluated.

**Table 1.**
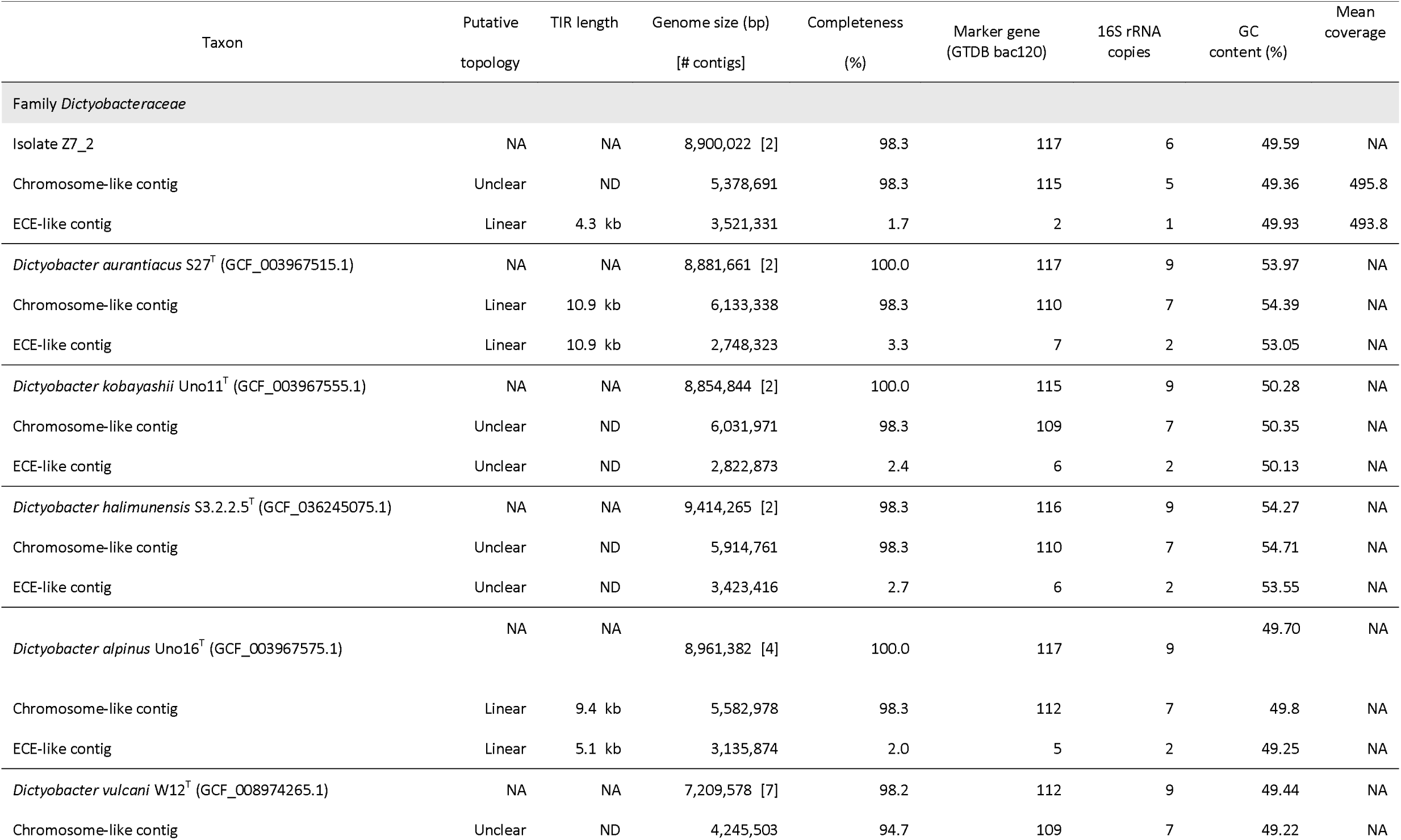
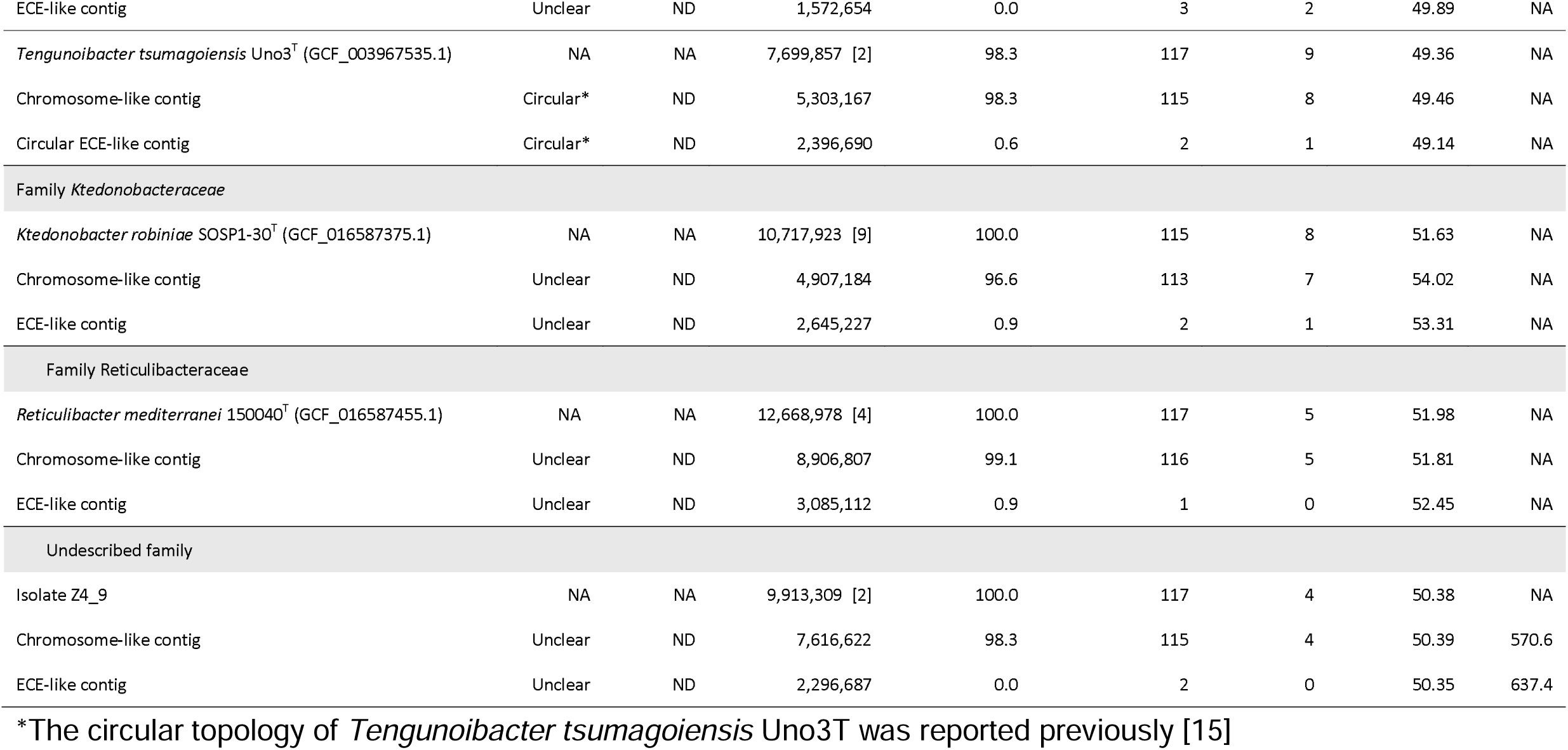
Summary of chromosome- and ECE-like contigs, genome features, and marker gene distribution in newly isolated and representative *Ktedonobacteria* genomes. This table summarizes the presence of putative extrachromosomal element (ECE)-like contigs in two newly isolated strains from Mt. Zao and selected genomes of the class *Ktedonobacteria*. ECE-like contigs were defined as ≥1 Mbp contigs whose CheckM-based completeness was at least five times lower than the completeness expected from their size relative to the total genome. CheckM-based completeness was assessed for each contig of the selected genomes. “Putative topology” indicates whether the contig was inferred to be circular, linear, or unclear, based on terminal inverted repeats (TIRs) or assembly structure; ND = not determined, NA = not applicable. TIR length refers to the approximate size of terminal inverted repeats, where detected. Genome sizes are shown together with the total number of contigs in parentheses (see Supplementary Data 11 for details of small contigs). Mean Illumina coverage was calculated only for the two newly isolated strains Z7_2 and Z4_9.

Most ECE-like contigs encoded only 1–7 GTDB bac120 marker genes and often harboured 16S rRNA genes. Cumulative GC-skew (Z-curve) profiles showed distinct V-or inverted V-shaped extrema on CHR- and ECE-like contigs, consistent with putative replication-origin (ori) candidates in all strains except Z4_9 (Fig. 4A); the major V coincided with the ori predicted by Ori-Finder 2022. In CHR-like contigs, dnaA/dnaN (replication initiation) were colocalised with parA/parB (partitioning) as a contiguous locus. In contrast, Pfam HMM searches detected no canonical Rep-family initiators on ECE-like contigs; parA homologues were present on all ECE-like contigs except those from Z4_9 and, when present, localised near the putative ori peak, whereas parB was not detected on any ECE-like contig (Fig. 4A). Consistent with distinct maintenance systems, the obtained ParA/RepA phylogeny separated CHR- and ECE-derived homologues into two well-supported groupings; most ECE-like contig ParA sequences formed a coherent clade, whereas ParA from Z7_2 and *Ktedonobacter robiniae* branched independently (Supplementary Fig. S4).

**Fig. 4.**
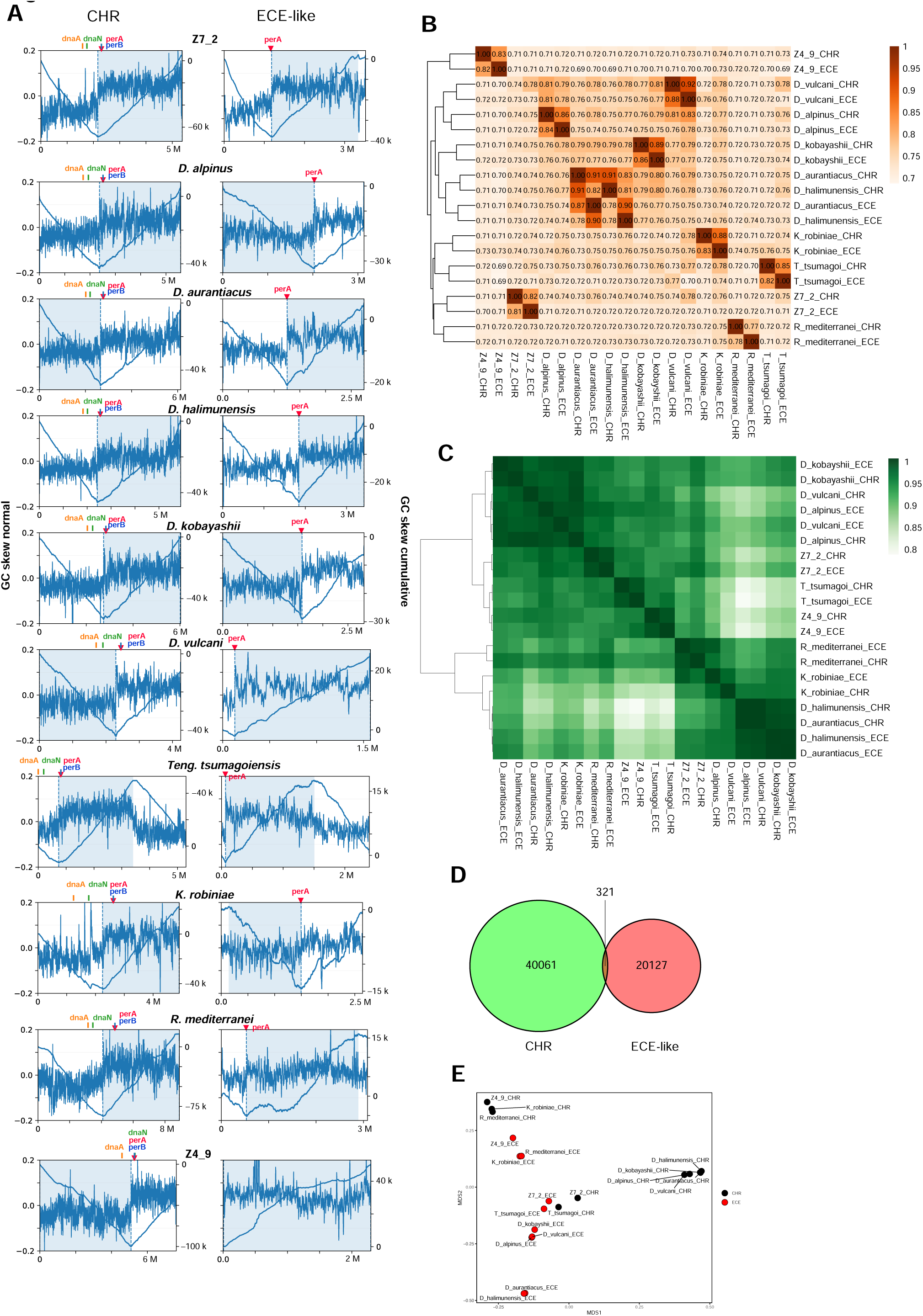
Genomic and compositional distinctions between CHR- and ECE-like contigs in the class *Ktedonobacteria*. **(A)** Cumulative GC skew profiles of CHR- and ECE-like contigs. Paired plots of normalised GC skew (blue line, left y-axis) and cumulative GC skew (shaded curve, right y-axis) are shown for each representative CHR–ECE pair. Gene markers for *dnaA*, *dnaN*, *parA*, and *parB* are indicated above each plot. The ori, predicted using Ori-Finder 2022, is shown by vertical dashed lines. The x-axis represents genomic coordinates (Mb), and the right y-axis for cumulative GC skew is scaled in arbitrary units (k). **(B)** Heatmap of ANI among CHR- and ECE-like contigs, calculated using PyANI (ANIb). Values indicate pairwise ANI and are shown by colour intensity (and numbers). Host–ECE pairs are highlighted with black circles. **(C)** Heatmap of codon-usage correlation (Spearman’s rank) among CHR- and ECE-like contigs. **(D)** Venn diagram showing the number of orthologous genes shared between CHR- and ECE-like contigs and those detected exclusively in each group. **(E)** Comparison of CHR- and ECE-like contigs based on gene-content variation. PCoA was performed for gene presence/absence patterns across 7,586 orthologous gene clusters (identified using Panaroo in strict mode). Pairwise dissimilarities were calculated using Jaccard distance. Each point represents a CHR- or ECE-like contig; filled and open symbols indicate CHR- and ECE-like replicons, respectively. PERMANOVA confirmed a statistically significant separation between the two groups (pseudo-F = 2.04, R² = 0.10, p = 0.004). ori, origin of replication; ANI, average nucleotide identity; PCoA, principal coordinate analysis; ECE, extrachromosomal element; CHR, chromosome.

Although the GC content of the ECE-like contig was nearly identical to that of the host CHR in each strain (Table 1), the replicons were nucleotide-dissimilar (fastANI 77–92% (Fig. 4B); Mash 0.22–0.26 based on 15-mers (Supplementary Data S12; Fig. 4B)). Despite this divergence, ECE-like codon usage correlated most strongly with the host CHR, and the two replicons often clustered together (Fig. 4C). In *D. halimunensis* and *D. aurantiacus*, the CHR-like and ECE-like contigs of each strain formed a pair, consistent with ANI-based clustering. For Z7_2 and Z4_9, mean Illumina coverage was ∼1:1 for CHR- and ECE-like contigs (Table 1).

Pan-genome analysis in Panaroo identified 40,061 genes unique to CHR-like contigs, 20,127 genes unique to ECE-like contigs, and only 321 shared genes (Fig. 4D), indicating strong compositional partitioning. Orthologue sharing among ECE-like contigs was sparse and largely strain-specific (Supplementary Data S13). Jaccard-based PCoA separated CHR- and ECE-like contigs into distinct clusters (Fig. 4E), which was supported by PERMANOVA results (pseudo-F = 2.04, R² = 0.10, p = 0.004).

Functionally, CHR-like contigs were enriched in housekeeping processes, whereas ECE-like contigs were enriched in transport/metabolism, secondary metabolism, and mobile functions (COG P, G, Q, K, and L) (Supplementary Fig. S5). Taken together, these ECE-like contigs share key chromid-like properties (e.g. large size, GC content and sequencing coverage similar to the primary CHR, host-correlated codon usage, and ParA-based maintenance signatures), which distinguish them from ordinary plasmids.

### BGC enrichment and associations with mobility genes in ECE-like contigs with cross-replicon homology

The genome maps of ten strains were annotated for BGCs, mobility-associated genes (integrases, recombinases, and transposases), and high-identity segments shared between CHR- and ECE-like contigs (≥98% nucleotide identity over ≥300 bp) (Fig. 5). Each genome encoded 25–238 predicted mobility-associated genes distributed across both replicons, and in several cases, the shared high-identity segments overlapped these loci (Fig. 5). Putative terminal inverted repeats (TIRs; ≥5 kb, ≥99% identity), often considered hallmarks of linear replicons, were detected on CHR- and ECE-like contigs in *D. aurantiacus* and *D. alpinus*, and only on the ECE-like contigs in strain Z7_2.

**Fig. 5.**
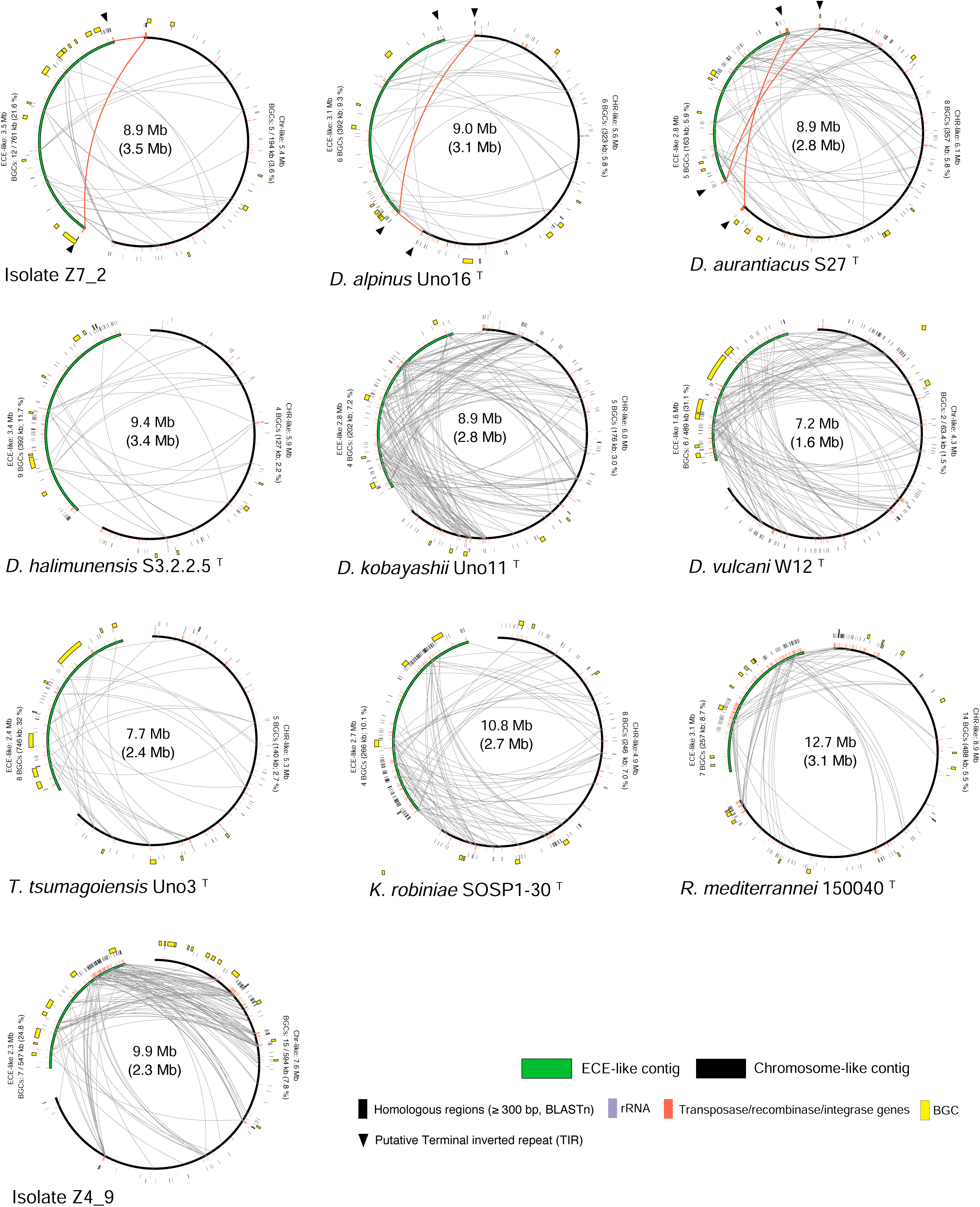
Shared homologous regions and distribution of BGCs and mobility genes on CHR- and ECE-like contigs. Circular maps are shown for ten strains. The CHR- and ECE-like contigs are indicated in black and green, respectively. Grey ribbons connect pairwise homologous regions detected by BLASTn (identity ≥ 98%, length ≥ 300 bp). Orange ribbons indicate long, high-identity homologous segments (length ≥ 5 kb, identity ≥ 99%). Black triangles denote putative TIRs. Gene features are overlaid as follows: yellow boxes, BGCs (predicted with antiSMASH v7.1.0); red ticks, mobility-associated genes (integrases, recombinases, and transposases); purple ticks, rRNA operons. The centre of each panel shows the total length of the CHR–ECE pair, with the ECE-like contig length in parentheses. For each replicon, ring-side labels report its length and BGC burden in the format “8 BGCs (746 kb; 32%)”. Each percentage indicates the fraction of the corresponding replicon (CHR- or ECE-like contig) covered by BGCs. TIR, terminal inverted repeat; ECE, extrachromosomal element; BGC, biosynthetic gene cluster; CHR, chromosome.

Consistent with this topology, cross-replicon homology blocks with ≥5 kb and ≥ 99% identity were mapped to the respective termini in all three strains, a pattern consistent with recent or ongoing recombination or segmental exchange between CHR- and ECE-like contigs.

BGCs occurred on both replicons but were generally enriched on ECE-like contigs. The fraction of each ECE-like contig sequence carrying BGCs exceeded that of the corresponding CHR-like contig in all strains examined and was particularly large in *T. tsumagoiensis* (32.0%), *D. vulcani* (31.1%), Z4_9 (24.8%), and Z7_2 (21.6%) (Fig. 5).

Length-corrected binomial tests and within-replicon shuffling were used to quantify spatial associations between mobility loci and BGCs (Supplementary Fig. 6A–C). Across replicons, mobility loci were depleted within BGCs (FE_within = 0.78; binomial p = 0.0023) but enriched near BGC boundaries, with the strongest enrichment observed in short flanks (L = 2 kb: FE_flank = 1.77; binomial p = 0.0099; permutation p = 0.022).

Overall, these analyses indicate that mobility loci preferentially occurred near BGC boundaries rather than inside BGCs. Distance-to-boundary analyses similarly showed enrichment of mobility loci within 0–2 kb of BGC boundaries (1.63x; permutation p = 0.022) (Supplementary Fig. S6C).

### Differences in BGC subclass composition between CHR- and ECE-like contigs

Principal coordinate analysis of subclass-level BGC profiles separated CHR- and ECE-like contigs (Fig. 6A), which was confirmed by PERMANOVA results (pseudo-F = 3.42, R² = 0.16, p = 0.001), indicating a replicon-type structuring of BGC repertoires.

**Fig. 6.**
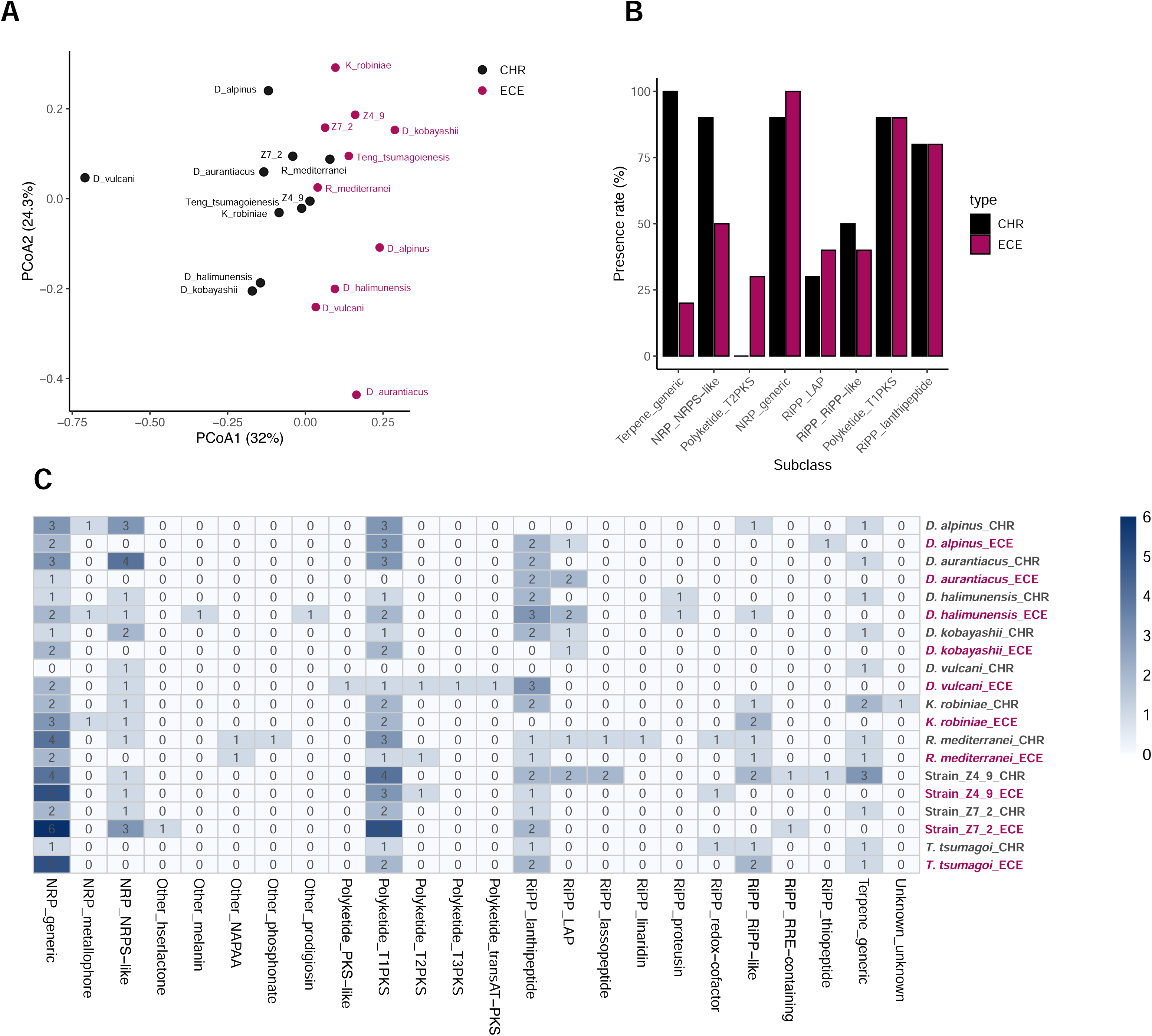
Subclass-level distribution of biosynthetic gene clusters on CHR- and ECE-like contigs in *Ktedonobacteria*. **(A)** PCoA of Bray–Curtis dissimilarities calculated from subclass-level BGC relative abundances. Points are differently coloured based on replicon type (CHR-like contig, black; ECE-like contig, red) and labelled with genome identifiers. PERMANOVA results showed significant separation by replicon type (pseudo-F = 3.42, R² = 0.16, p = 0.001). **(B)** Bar plot showing presence rates (%) of BGC subclasses in CHR- (black) and ECE-like contigs (red). Only subclasses detected in at least three genomes were included. No subclasses were significant after multiple-testing correction (exact McNemar test with Benjamini–Hochberg adjustment; p ≥ 0.05). **(C)** Heatmap showing the counts of GCFs assigned to each BGC subclass across replicons. Rows and columns correspond to individual replicons and BGC subclasses, respectively. Cell shading indicates the number of detected GCFs per subclass, with darker shades indicating higher counts. GCF, gene-cluster family; BGC, biosynthetic gene cluster; PCoA, principal coordinate analysis; ECE, extrachromosomal element; CHR, chromosome.

Subclass frequency comparisons showed no significant differences after multiple-testing correction (Fig. 6B), but terpene clusters were largely confined to CHR-like contigs, whereas type II PKS clusters were detected only on ECE-like contigs in our dataset.

Heatmap visualisation showed genome-level replicon signatures, with several BGC subclasses encoded exclusively on ECE-like contigs (Fig. 6C). For example, in Z7_2, a homoserine-lactone (hserlactone) cluster was confined to the ECE-like contig, and in *D. halimunensis*, multiple subclasses (including RiPP-related and pigment-associated clusters) occurred only on ECE-like contigs.

## Discussion

*Ktedonobacteria* is a morphologically distinctive lineage within the phylum *Chloroflexota* [15, 45] and is often abundant in extreme terrestrial environments, including volcanic soils and polar regions (Fig. 1A) [9, 10, 13, 46]. Our analyses indicated that this bacterial class encodes a rich and compositionally novel repertoire of BGCs, many of which are located on large chromid-like ECEs. The isolated representative of the previously uncultivated, BGC-enriched clade K1, together with the *Dictyobacter* isolate described here, provides a basis for experimental validation and future exploration of their secondary metabolites.

Overall, 25% of the 183 dereplicated ktedonobacterial genomes examined were shown to encode ≥10 nonredundant GCFs each (Fig. 1B), indicating that BGC richness is shared across multiple families (Fig. 1A) rather than restricted to a few taxa. This trait was observed across several family-level clades (e.g. K1, K12, K13, and RTK), consistent with phylogenetically widespread biosynthetic potential. Novelty analyses characterised *Ktedonobacteria* as a class comparable to canonical discovery lineages: against the BiG-FAM dataset (1.225 million predicted BGCs)[35], 52.6% of ktedonobacterial BGCs exceeded the novelty threshold (Euclidean distance > 900) [38] versus 44.9% for *Streptomyces* (Fig. 2B(i)); whereas against the MIBiG dataset [47], 96.0% exceeded the novelty cutoff (d > 0.5) [48] (Fig. 2B(ii)). Clustering analysis further indicated low intra-genomic redundancy, with an approximately one-to-one correspondence between BGCs and GCFs per genome, contrasting with the results obtained for *Streptomyces* (Fig. 2C). Overall, these patterns suggest that individual ktedonobacterial genomes encode compact, structurally diverse BGC repertoires, which may benefit discovery efforts that prioritise breadth over closely related chemotypes. An important caveat is that these estimates are likely conservative.

Cultured genomes harboured roughly twice as many GCFs as MAGs on average, suggesting that chromid-like secondary replicons are underrepresented in short-read MAG surveys due to marker-based filtering and misbinning.

A salient ecological question is why *Ktedonobacteria* carry such abundant and diverse BGCs. Secondary metabolism appears to be associated with morphological complexity [49]: the K1 strain Z4_9 and the *Dictyobacter* strain Z7_2 harbour numerous distinct GCFs and exhibit complex filamentous morphologies (Fig. 1A, B). Strain Z4_9 belongs to a deeply branching, previously uncultivated clade that likely represents a novel, BGC-enriched family, whereas Z7_2 expands cultivation-based diversity within *Dictyobacteraceae*. Filamentous, actinomycete-like morphologies were widely observed across *Ktedonobacteria* (Fig. 1A), raising the possibility of a functional coupling between secondary metabolism and morphological differentiation that assists in surface-associated growth and interactions within microbial communities. The linkage between these traits is well documented in *Actinomycetes* [49], and *Ktedonobacteria* may have evolved an analogous strategy by integrating morphological differentiation with chemically mediated defence. The isolates examined in this study provide an opportunity for direct tests linking genome architecture, morphology, ecological function, and secondary metabolites.

BGC distributions differed across phylogenetic ranks (family and order; Fig. 3A, Supplementary Fig. S3), and an association was also detected between ecological context and secondary-metabolite potential (Fig. 3B, Supplementary Fig. S3).

Specifically, the latter is likely shaped by shared ancestry, a hypothesis consistent with the preferential vertical preservation of core biosynthetic pathways [50]. GLMM analyses aiming to separate habitat effects from phylogenetic signal identified genomes with enriched RiPP_redox-cofactor clusters from samples obtained from volcanic or other stress-prone soils (Fig. 3B). This pattern was consistent with a possible association between RiPP_redox-cofactor loci and oxidative or nutrient-limited environments [44, 51]. It remains to be determined whether these loci are expressed and what products (if any) they yield.

Another key question is why BGCs may preferentially accumulate on ECE-like contigs. Representative cultured strains from several ktedonobacterial families have been shown to harbour 1.6–3.5 Mb secondary replicons with properties distinct from those of typical small plasmids [22]. In each strain examined in the present study, the GC content and sequencing coverage of the ECE-like contig were nearly identical to the primary CHR, and yet the former showed only 77%–92% ANI and substantial gene-content divergence (Fig. 4A, B, D; Table 1). Codon-usage patterns of the two replicons were strongly correlated and often clustered together (Fig. 4C), and the obtained ParA/RepA phylogeny recovered distinct but coherent clades for CHR- versus ECE-derived homologues (Supplementary Fig. S4). Functional profiles support this separation. CHR-like contigs were enriched in housekeeping processes, whereas

ECE-like contigs were enriched in transport, secondary metabolism, and mobility functions (Supplementary Fig. S5). Collectively, these observations indicate that ECE-like contigs are large, host-adapted secondary replicons with chromid-like features. They share nucleotide composition and translational signatures with the primary CHR [22, 24] and yet carry a distinct, functionally specialised gene repertoire [22]. Although their evolutionary origin remains uncertain, these host-associated signatures and the enrichment in adaptive functions provide a plausible context for BGC accumulation on ECE-like replicons.

The ecological and evolutionary roles of chromids remain a subject of debate, but a widely discussed view is that they allow adaptive, environmentally responsive genes to accumulate on a secondary replicon while retaining core housekeeping functions on the primary CHR [22, 24, 52]. If the ECE-like replicons in *Ktedonobacteria* fulfil a similar role, their large size and bias towards transport, secondary metabolism, and mobility functions would be consistent with such a division of labour and may contribute to the relatively large genome sizes observed in this bacterial class [15].

Our analyses indicate that mobility loci are depleted within BGCs but enriched near BGC boundaries, consistent with preferential insertion/rearrangement at cluster edges and potential reshuffling of biosynthetic modules while maintaining internal cluster integrity.

Extensive homologous segments are shared between CHR- and ECE-like contigs, and genes related to environmental adaptation are concentrated on ECE-like replicons (Fig. 5). These patterns are consistent with mobile-element activity and homologous recombination facilitating the exchange of environmentally responsive genes, including BGCs, between replicons [53]. One possible scenario is that ECE-like replicons undergo a comparatively rapid evolutionary change [22, 54], sampling and reshuffling adaptive loci, some of which may later become stabilised on the primary CHR, implying a functional division of labour between replicons.

Consistent with this interpretation, terpenes involved in carotenoid biosynthesis were broadly detected on CHR-like contigs (Fig. 6B), consistent with their conserved roles in light and oxidative stress responses [55]. In contrast, ECE-like contigs frequently encoded BGC subclasses absent from the corresponding CHR (Fig. 6C). This pattern was particularly evident in *D. halimunensis*, whose ECE-like contig harboured RiPP_RiPP-like, prodigiosin, melanin, and metallophore clusters absent from the CHR (Fig. 6C). In environments subjected to fluctuations or disturbance, such ECE-restricted modules might provide flexible, niche-dependent functions that can be gained or lost over short evolutionary timescales [22].

Terminal homology between CHR- and ECE-like contigs was detected in a subset of ktedonobacterial genomes, consistent with occasional homologous recombination.

In summary, by integrating newly cultivated representatives with metagenome-derived genomes, we provide a genome-wide assessment of secondary metabolism and replicon architecture in *Ktedonobacteria*. Our analyses indicate clade-wide enrichment of diverse and apparently novel BGCs and reveal the pervasive presence of chromid-like ECEs that preferentially accumulate BGCs and exhibit mobility-associated functions. These findings establish a foundation for future research to validate ECE autonomy and maintenance and to functionally characterise ECE-borne BGCs through expanded cultivation and targeted experiments.

## Acknowledgements

We thank our laboratory staff and relevant authorities for their support with sequencing and sampling.

## Author contributions

S.Y. conceived and designed the study, performed experiments and data analyses, and drafted the manuscript. Z.Y. performed bioinformatic analyses, including structure-based predictions supporting biosynthetic product inference. S.T. provided expert input on secondary metabolism and critically reviewed the logical structure of the manuscript. C.Y. contributed to TOC measurements and interpretation. Y.N. contributed to next-generation sequencing and data processing. M.S. and K.T. contributed to SEM analyses and interpretation. S.Yam. and N.O. assisted with field sampling and sample processing. M.K.R., F.N., and W.S. provided strain-specific expertise on *D. halimunensis* and contributed to interpretation. Y.I. supervised the project and provided resources. All authors critically revised the manuscript, approved the final version, and agree to be accountable for all aspects of the work.

## Data availability

Shotgun metagenome reads are available in the DDBJ Sequence Read Archive (DRA) under BioProject PRJDB35689 (runs DRR728410–DRR728417). Raw isolate genome reads (Z7_2 and Z4_9) are available under the same BioProject PRJDB35689 (runs DRR905084–DRR905085 and DRR905086–DRR905087). Metagenome-assembled genome (MAG) assemblies generated in this study have been deposited in DDBJ/ENA/GenBank, and accession numbers are currently under processing. Complete genome assemblies of strains Z7_2 and Z4_9 have also been submitted and are currently under processing. The 16S rRNA gene sequences of these isolates are available under LC811933 (Z7_2) and LC811932 (Z4_9). Source data for Figs. 2–4 and 6 and Supplementary Fig. 4 are provided in the Supplementary Data. Additional information is available from the corresponding author upon reasonable request.

