## Supplementary Figures for "Chromid-like secondary replicons as key sites of biosynthetic gene clusters in *Ktedonobacteria*"

**This PDF includes;**

Supplementary Figure 1 to Supplementary Figure 6

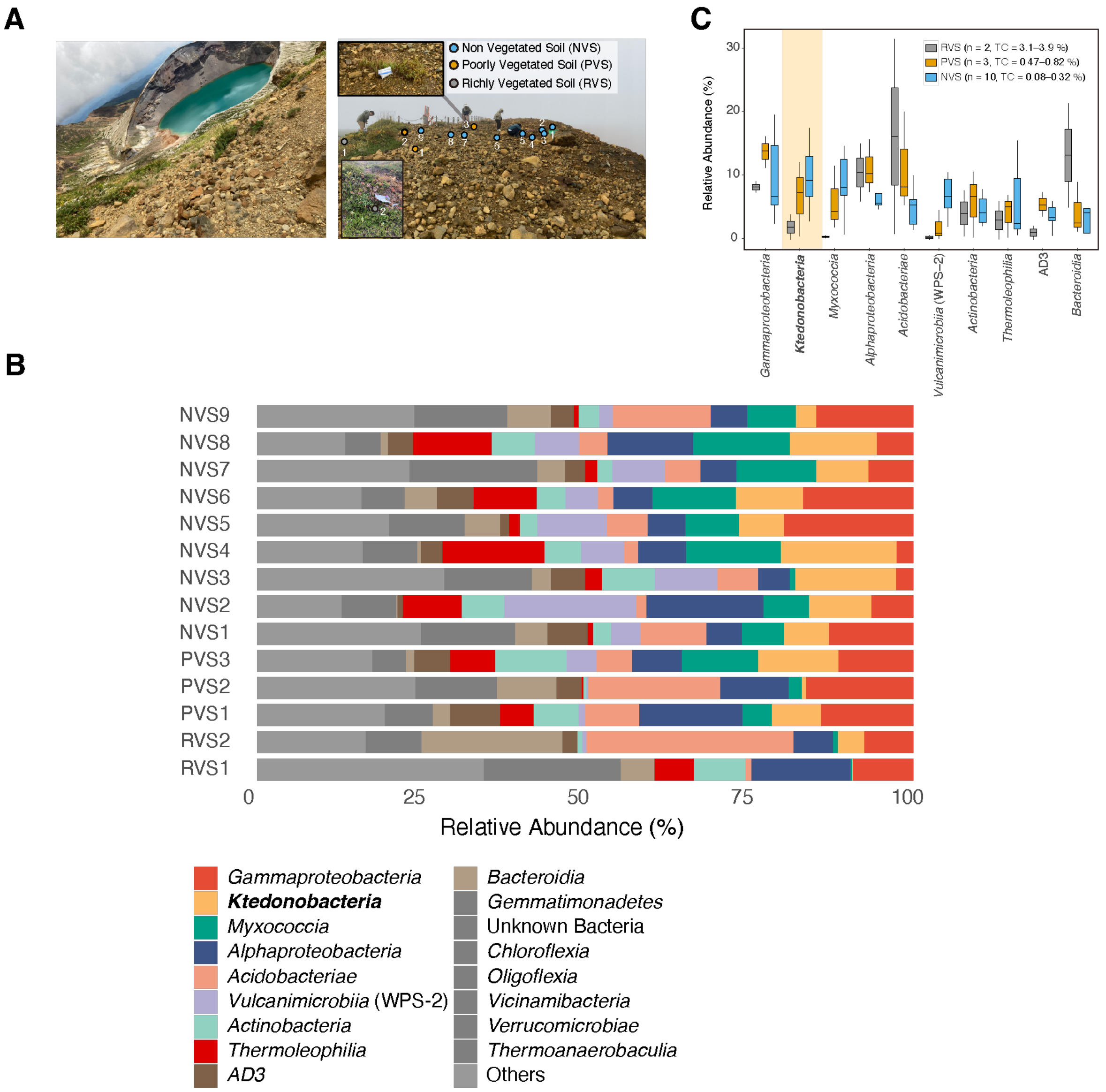


**Supplementary Figure 1. Sampling design and class-level bacterial composition in soils from Zao Volcano.**

**(A)** Sampling sites. Left: panoramic view of the crater lake Okama at Mt. Zao, Japan. Right: close-up of the sampling area, showing non-vegetated soils (NVS), poorly vegetated soils (PVS), and richly vegetated soils (RVS). Sampling points are color-coded according to vegetation cover and associated total organic carbon (TOC) content: RVS (n = 2, TOC = 2.04%), PVS (n = 3, TOC = 0.46–0.99%), and NVS (n = 10, TOC = 0.25–0.41%).

**(B)** Class-level bacterial community composition in individual soil samples. Stacked bar plots show the relative abundance of bacterial classes across NVS, PVS, and RVS. Taxa with >5% relative abundance in at least one sample are shown individually; all others are grouped as "Others." Taxonomic assignment is based on 16S rRNA gene sequences.

**(C)** Relative abundances of the ten most abundant bacterial classes in NVS and PVS soils, based on 16S rRNA gene (V4 region) amplicon sequencing. Boxes indicate interquartile ranges with medians.

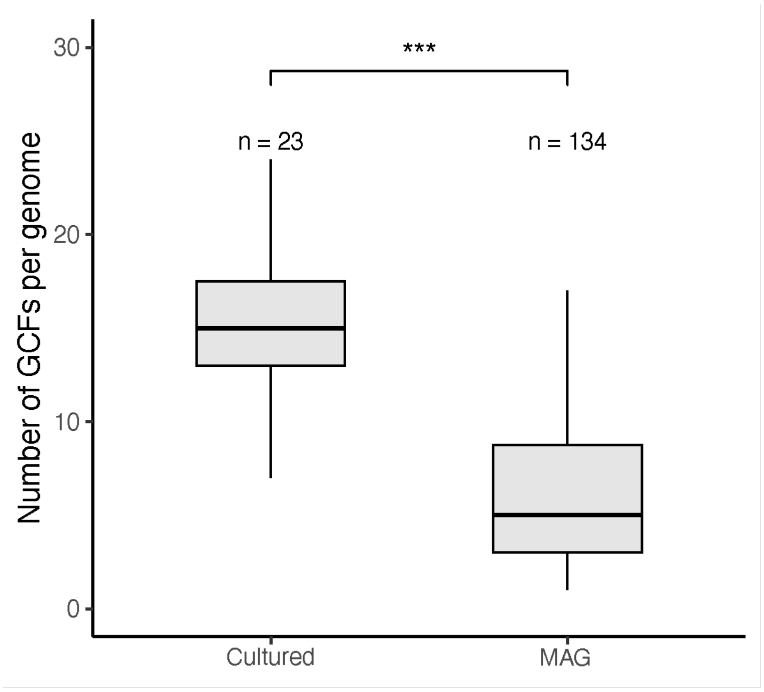


**Supplementary Fig. 2 Comparison of the number of GCFs per genome between cultured isolates and MAGs of *Ktedonobacteria*.**

The number of GCFs per genome was plotted for each group. Wilcoxon rank-sum test was used to evaluate statistical significance (*p < 0.05, **p < 0.01, ***p < 0.001). Sample sizes are indicated below each group

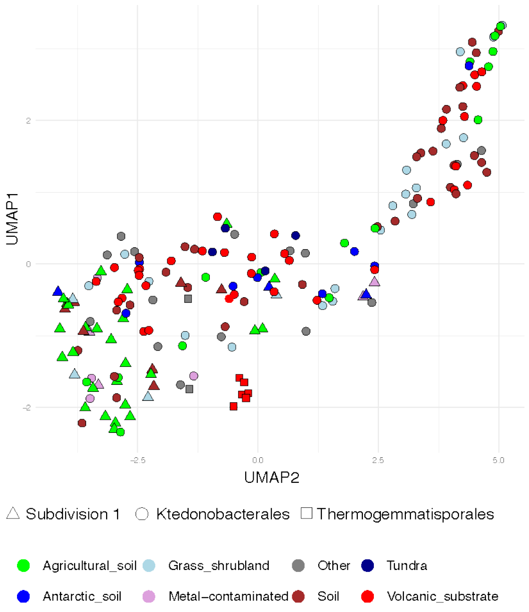


**Supplementary Fig. 3 UMAP projection of GCF subclass profiles across genomes, grouped by taxonomic and environmental attributes.**

Each point represents a genome, plotted based on its secondary metabolite subclass composition derived from BiG-SLiCE classification. Subclass names are shown on the X-axis in a “Class_Subclass” format (e.g., RiPP_bottromycin). Shapes indicate major taxonomic groups (Clade S, Thermogemmatisporaceae, and others), and colors denote ecosystem of origin (e.g., volcanic substrate, tundra, soil). PERMANOVA showed significant clustering by taxonomic group (family: $p = 0.001$; order: $p = 0.001$) and environmental source ($p = 0.011$).

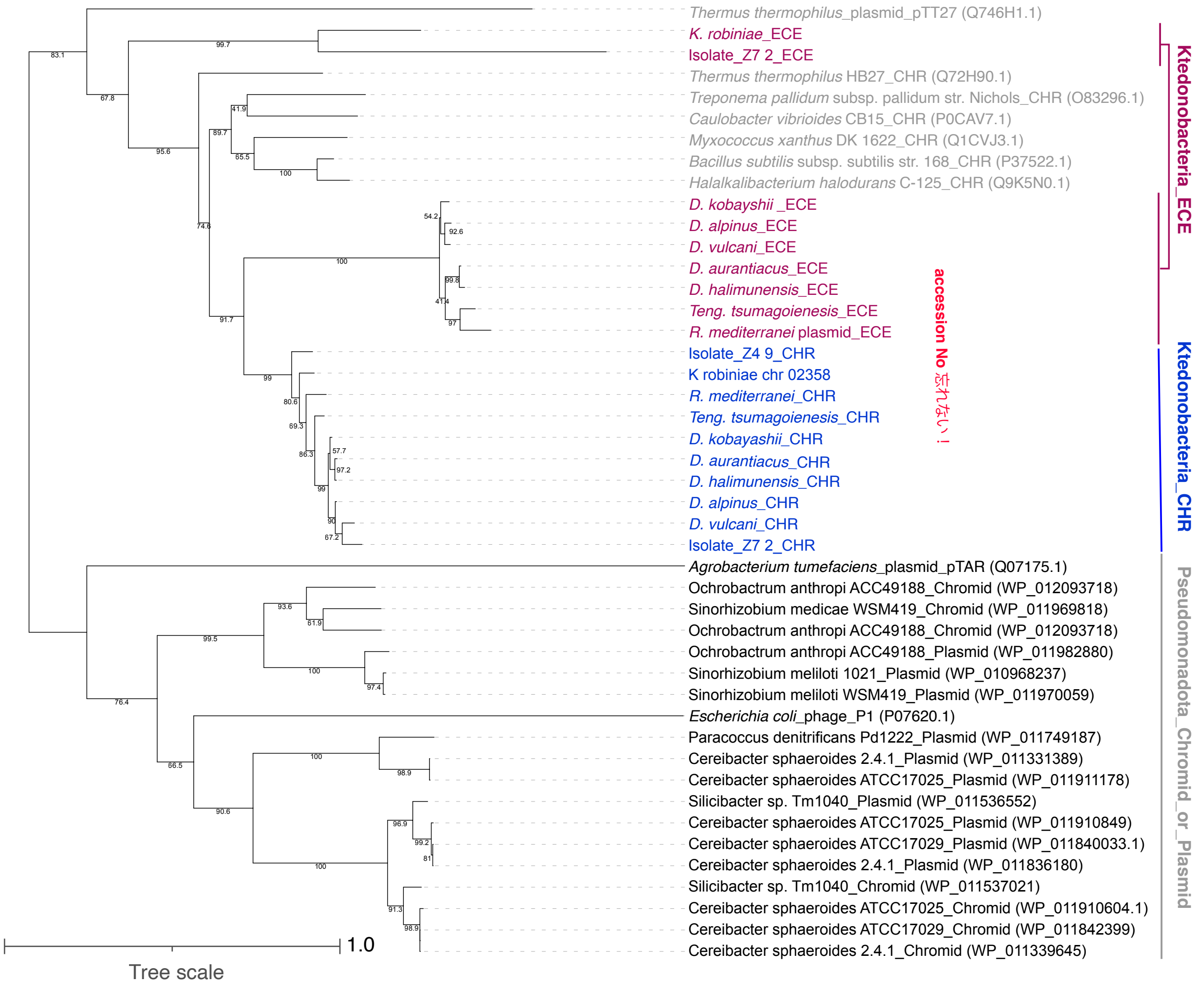


**Supplementary Fig. 4 Maximum-likelihood phylogeny of ParA/RepA homologs from representative CHR- and ECE-like contigs.**

Protein sequences were aligned with MUSCLE v3.8.425, trimmed with trimAl (gap threshold = 0.5, similarity threshold = 60), and analyzed in IQ-TREE v2.3.6 under the Q.pfam+I+G4 substitution model. Branch support values are shown as SH-aLRT (left) and ultrafast bootstrap (right) percentages (both with 1,000 replicates).

*Ktedonobacteria* sequences are highlighted in red (ECE-like contigs-derived) and blue (CHR-like contigs), while other representative taxa are shown in black.

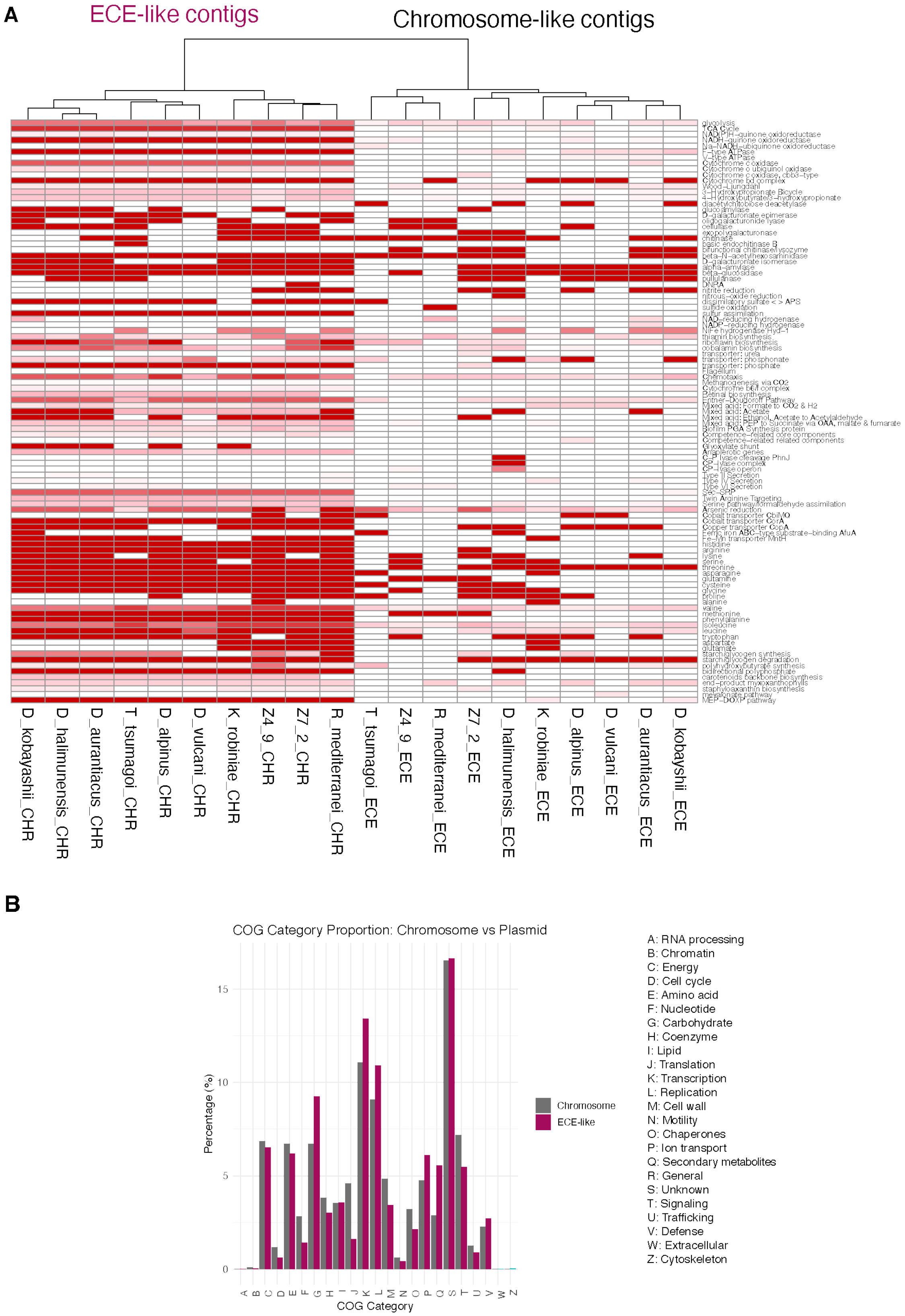


**Supplementary Fig. 5 Functional partitioning between CHR-like and ECE-like contigs in *Ktedonobacteria*.**

**(A)** Heatmap of KEGG functional categories (rows) across contigs (columns) from the isolates shown. Columns are hierarchically clustered and grouped by replicon type: ECE-like and CHR-like contigs. Cell color indicates the normalized representation (0–1) of genes assigned to each category within a contig. Functional annotation used eggNOG-mapper v2.1.12 (eggNOG v5.0.2), and KEGG terms were parsed from its output.

**(B)** Relative proportions of COG one-letter functional categories on CHR-like (grey) and ECE-like contigs (magenta) computed from the same annotations; category labels (A–Z) are listed at right.

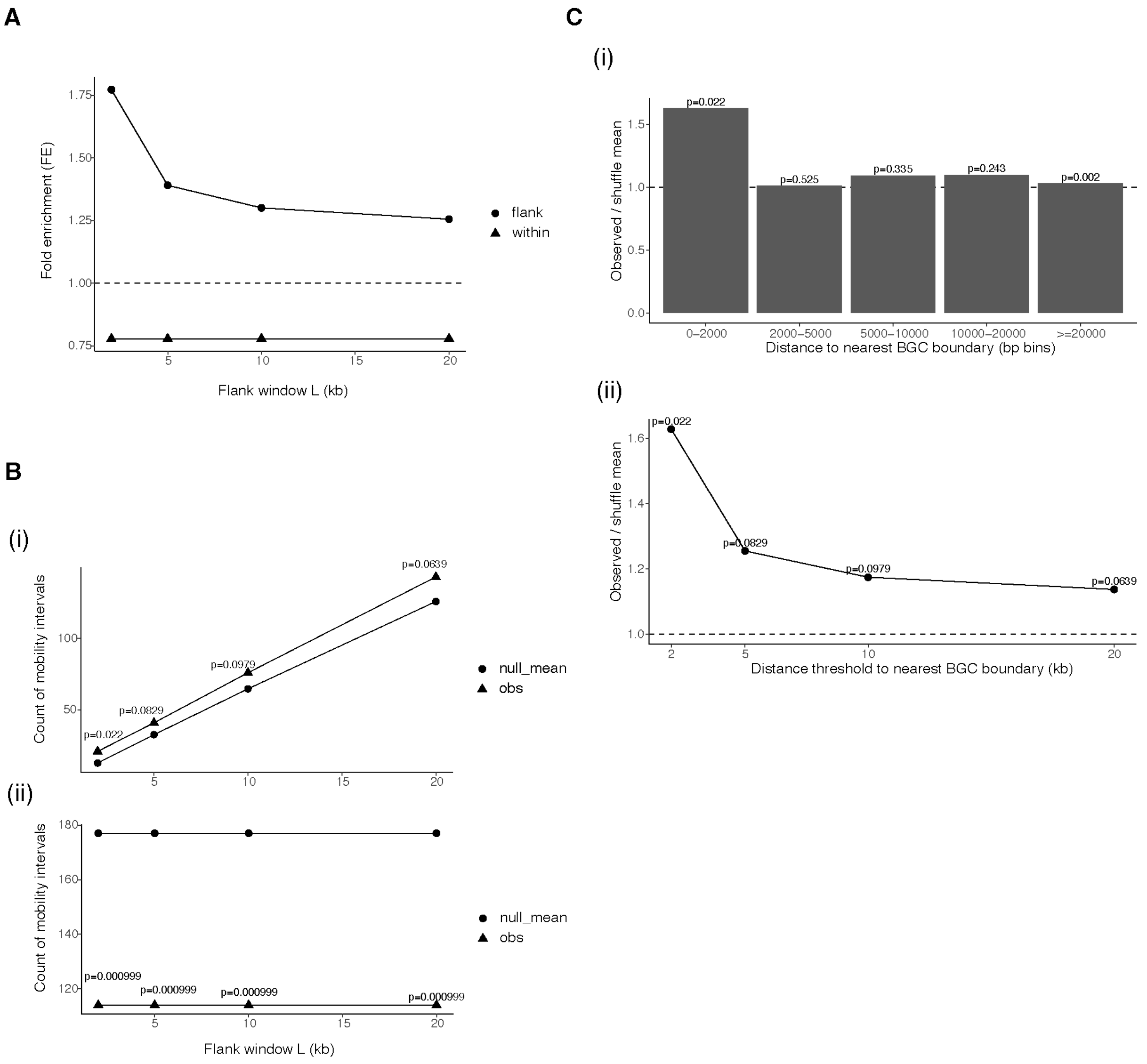


**Supplementary Fig. 6 Spatial association between mobility loci and biosynthetic gene clusters**

Spatial relationships between mobility loci and biosynthetic gene clusters (BGCs) assessed using length-corrected enrichment analyses and contig-restricted permutation tests.

**(a)** Fold enrichment (FE) of mobility loci within BGC bodies (within; triangles) and in flanking regions surrounding BGCs (flank; circles) across different flank window sizes (*L* = 2, 5, 10, and 20 kb).

Fold enrichment was calculated as the ratio of the observed fraction of mobility loci in a given region to the fraction expected under a random distribution proportional to region length.

The dashed horizontal line indicates the expectation under random distribution (FE = 1).

**(b)** Robustness of the enrichment patterns evaluated by contig-restricted shuffling (1000 permutations), in which mobility intervals were randomly repositioned within each replicon while preserving interval lengths.

(i) Observed counts of mobility loci in BGC flanking regions (triangles) compared with the

mean expected counts from shuffling (circles).
(ii) Observed counts of mobility loci overlapping BGC bodies (triangles) compared with shuffle-based expectations (circles).
Empirical one-sided permutation *P* values are shown for each window size.
**(c)** Distance-based analysis of mobility loci relative to BGC boundaries. Mobility loci overlapping BGC bodies were excluded from this analysis.
(i) Enrichment in discrete distance bins, shown as the ratio of observed counts to the shuffle mean for mobility loci located at increasing distances from the nearest BGC boundary.
(ii) Cumulative enrichment of mobility loci located within the indicated distance thresholds from the nearest BGC boundary.
The dashed horizontal lines in panels (b) and (c) indicate expectations under the shuffle-based null model (observed/shuffle mean = 1).
Together, these analyses show that mobility loci are depleted within BGC bodies but preferentially accumulate in the immediate vicinity of BGC boundaries, with the strongest enrichment observed within approximately 2 kb of BGC borders.
