## Supplementary_Methods for "Chromid-like secondary replicons as key sites of biosynthetic gene clusters in *Ktedonobacteria*"

Affiliations:
1, RIKEN Center for Sustainable Resource Science, 3-1-1 Koyadai, Tsukuba, Ibaraki, 305-0074, Japan
2, RIKEN Center for Sustainable Resource Science, 2-1 Hirosawa, Wako, Saitama, 351-0198, Japan
3, Department of Biology, Faculty of Mathematics and Natural Sciences, Universitas Indonesia, Kampus UI Depok, 16424, Indonesia
4, Center of Excellence for Indigenous Biological Resources-Genome Studies, Faculty of Mathematics and Natural Sciences, Universitas Indonesia, Kampus UI Depok, 16424, Indonesia
2, RIKEN Center for Sustainable Resource Science, 1-7-22 Suehiro, Tsurumi, Yokohama, Kanagawa, 230-0045, Japan

Corresponding authors:
Author 1: Dr Shuhei Yabe,
Author 2: Dr Yasunori Ichihashi,

**Method S1. Soil sampling and TOC measurement**

Soil samples were collected on 29 September 2023 from volcanic barren soils on the crater rim of Lake Okama, Mount Zao, Miyagi, Japan (38.1321°N, 140.4468°E; 1,680 m a.s.l.; Supplementary Figure 1a). Fifteen surface soils (0–10 cm) were obtained using sterile spatulas and gloves, placed in sterile tubes or zipper bags, and transported on ice to the laboratory within 48 h. Based on vegetation cover and total organic carbon (TOC) content, the samples were classified as non-vegetated soils (NVS; n = 10, 0.25–0.41% TOC), poorly vegetated soils (PVS; n = 3, 0.46–0.99% TOC), and richly vegetated soils (RVS; n = 2, 2.04% TOC). Subsamples were stored at –20 °C for DNA extraction, at 4 °C or –80 °C for cultivation, and ~50 g was air-dried at 25 °C for TOC analysis.

TOC was quantified by treating 0.5 g of soil with 2 M HCl to remove inorganic carbon, followed by 12 h venting, freeze-drying, and measurement using multi EA 4000 (analytikjena, Jena, Germany). Measurements were performed in triplicate and reported as mean ± SD.

**Method S2. DNA extraction, sequencing, assembly, and MAG processing**

Freeze-dried soils (2.0 g) were cryogenically pulverized after liquid-nitrogen immersion in 3 mL tubes with a steel bead using a Multi-Beads Shocker at 3,000 rpm for 15 s (Yasui Kikai, Osaka, Japan). For 16S amplicons, DNA (90–500 mg soil) was extracted with the Extrap Soil DNA Kit Plus v2 (BioDynamics Laboratory, Tokyo, Japan); the V4 region was amplified with primers 515F/805R carrying Illumina overhangs and sequenced on a MiSeq (2 × 300 bp)[26]. Reads were processed in QIIME 2 v2024.10.1[2] with DADA2[3], and taxonomy was assigned using a naïve Bayes classifier trained on SILVA 138 (scikit-learn v1.4.2)[4].

For shotgun metagenomics, DNA from 0.5 g soil (NVS3: 10 g using ISOIL Large for Beads v2, Nippon Gene) was purified with the DNeasy PowerSoil Pro Kit (Qiagen, Germantown, MD, USA). Nextera XT libraries (Illumina, Inc., San Diego, CA, USA) were sequenced on a NovaSeq X (Illumina, Inc., San Diego, CA, USA) (2 × 150 bp; 54.1 Gbp total, mean 6.76 Gbp per sample). Eight datasets (five NVS, three PVS) were processed with MetaWRAP v1.3.2[5]: adapters were trimmed with Trim Galore v0.5.0; assemblies were generated with MEGAHIT v1.1.3[6]; contigs <500 bp were removed; quality was assessed with QUAST v5.0.2.[7].

Contigs ≥1 kb were binned with MetaBAT2[8], MaxBin2[9], and CONCOCT[10], and refined with MetaWRAP[5] bin_refinement. Bins meeting ≥50% completeness and ≤10% contamination (CheckM v1.0.12) [11]were retained as MAGs, reassembled with SPAdes v3.13.0[12], and re-evaluated with CheckM, yielding 166 non-redundant MAGs. MAGs were taxonomically assigned with GTDB-Tk v2.1.1 (GTDB R207)[13]; one (Merged_NVS_Bin65) was excluded after phylogenomic inspection, leaving 20 *Ktedonobacteria* MAGs. Dereplication at 95% ANI using dRep v3.5.0 (fastANI) [14]produced 90 species-level clusters; within each, the representative was the genome maximizing (completeness − 5×contamination; N50 as tiebreaker).

**Method S3. Isolation, genome sequencing, and SEM of *Ktedonobacteria***

Fresh soil was imprinted onto 1/10-strength R2A (SHIOTANI M.S., Osaka, Japan) solidified with 1.5% (w/v) gellan gum and 0.2% (w/v) $CaCl_2 \cdot 2H_2O$, acidified to pH 5.0 (HCl), and supplemented with 30 mg $L^{-1}$ sodium azide [15]. Plates were incubated aerobically at 30 °C in the dark for 60 days. Colonies forming tough, deeply penetrating substrate mycelia were picked as putative *Ktedonobacteria*, lysed in alkali buffer (25 mM NaOH, 0.2 mM $Na_2$-EDTA), heated at 95 °C for 30 min, neutralized with 40 mM Tris-HCl (pH 7.5), and used as PCR templates. Nearly full-length 16S rRNA genes were amplified with primers 27F/1492R[16] using TaKaRa Ex Taq, sequenced bidirectionally (Macrogen Japan, Kyoto, Japan), and identified via the EZBioCloud Identify Service[17]. Four of 56 *Ktedonobacteria* colonies were purified by repeated streaking and maintained on acidified R2A–gellan at 4 °C. Two phylogenetically distinct strains (Z7_2 and Z4_9) were selected for genome sequencing. Genomic DNA was extracted with a modified Puregene protocol (Qiagen, Germantown, MD, USA) including enzymatic lysis (achromopeptidase, lysozyme, proteinase K; overnight at 37 °C)[18]. Long reads were generated by Bioengineering Lab (PacBio Revio; Pacific Biosciences of California, Inc. Menlo Park, CA, USA) and short reads by Macrogen Japan (NovaSeq X; Illumina, Inc., San Diego, CA, USA ) (2 × 151 bp). Long reads were assembled with Flye v2.9.6-b1802[19], polished with Pilon v1.24[20] using Illumina read alignments, and circularized with Circlator v1.5.5[21], yielding complete circular genomes for both strains.

For SEM, colonies were fixed overnight with vapor from 2% osmium tetroxide solution at room temperature. Then samples were coated with osmium by osmium coater (HPC-1SW, Vacuum Devise), and observed using a Hitachi SU8220 field-emission SEM at an accelarating voltage of 3 kV. Previously unpublished SEM images of *K. robiniae*[22], *Tg. onikobensis*[23], and *D. aurantiacus*[24] were included for comparison in Fig. 1a.

**Method S4. Phylogenomics of *Ktedonobacteria***

From the NCBI Genome database (May 3, 2025; https://www.ncbi.nlm.nih.gov/datasets/), we retrieved 318 *Ktedonobacteria* genomes; after removing a duplicate (*Thermogemmatispora hazakensis* SK20-1), 317 unique genomes remained (21 cultured, 296 MAGs). Only the public MAGs were dereplicated at 95% ANI with dRep v3.5.0 (fastANI)[14], yielding 140 representatives. To retain local microdiversity, we then added 21 Mount Zao MAGs and two new isolates (Z7_2, Z4_9) without dereplication, giving a final dataset of 184 genomes. All genomes were processed with GTDB-Tk v2.1.1 (bac120)[13]. Marker genes were aligned with gtdbtk align, resulting in a concatenated amino-acid alignment of 35,833 sites. The alignment was trimmed with TrimAl v1.4[25] (-gt 0.5 -cons 60) and used to infer a maximum-likelihood tree in IQ-TREE v2.3.6 (LG+F+G4; 1,000 ultrafast bootstraps); trees were visualized in iTOL[26].

Family-level clades were delineated with TreeCluster v1.0.4[27] (“max_clade”, distance threshold

0.45), which preserved the monophyly of *Ktedonobacterales* and *Thermogemmatisporales* while increasing resolution. At this threshold, *Ktedonobacteraceae*, *Reticulibacteraceae*, and *Thermosporotrichaceae* formed a single clade ("Clade RKT"). Within *Ktedonobacterales* we designated Clades K1–K15 (plus three singletons); one clade was recovered within *Thermogemmatisporales*. A deeply branching lineage distinct from both orders was provisionally named "Subdivision 1," comprising Clades S1–S9 and eight singletons.

Of the 21 Mount Zao MAGs, 20 fell within *Ktedonobacteria* (*Dictyobacteraceae*; Clades K2, K4, K11; one *Ktedonobacterales* singleton; and Subdivision 1, Clade S4). One MAG (Merged_NVS_bin65) fell outside the class and was excluded from downstream analyses. Clade assignments were annotated on the tree and integrated with BGC profiles, cultivation metadata, and replicon architecture for subsequent comparative analyses.

**Method S5 Phylogenomics of *Ktedonobacteria***

From the NCBI Genome database (May 3, 2025; https://www.ncbi.nlm.nih.gov/datasets/), we retrieved 318 *Ktedonobacteria* genomes; after removing a duplicate (*Thermogemmatispora hazakensis* SK20-1), 317 unique genomes remained (21 cultured, 296 MAGs). Only the public MAGs were dereplicated at 95% ANI with dRep v3.5.0 (fastANI)[14], yielding 140 representatives. To retain local microdiversity, we then added 21 Mount Zao MAGs and two new isolates (Z7_2, Z4_9) without dereplication, giving a final dataset of 184 genomes. All genomes were processed with GTDB-Tk v2.1.1 (bac120)[13]. Marker genes were aligned with gtdbtk align, resulting in a concatenated amino-acid alignment of 35,833 sites. The alignment was trimmed with TrimAl v1.4[25] (-gt 0.5 -cons 60) and used to infer a maximum-likelihood tree in IQ-TREE v2.3.6 (LG+F+G4; 1,000 ultrafast bootstraps); trees were visualized in iTOL[26].

Family-level clades were delineated with TreeCluster v1.0.4[27] ("max_clade", distance threshold 0.45), which preserved the monophyly of *Ktedonobacterales* and *Thermogemmatisporales* while increasing resolution. At this threshold, *Ktedonobacteraceae*, *Reticulibacteraceae*, and *Thermosporotrichaceae* formed a single clade ("Clade RKT"). Within *Ktedonobacterales* we designated Clades K1–K15 (plus three singletons); one clade was recovered within *Thermogemmatisporales*. A deeply branching lineage distinct from both orders was provisionally named "Subdivision 1," comprising Clades S1–S9 and eight singletons.

Of the 21 Mount Zao MAGs, 20 fell within *Ktedonobacteria* (*Dictyobacteraceae*; Clades K2, K4, K11; one *Ktedonobacterales* singleton; and Subdivision 1, Clade S4). One MAG (Merged_NVS_bin65) fell outside the class and was excluded from downstream analyses. Clade assignments were annotated on the tree and integrated with BGC profiles, cultivation metadata, and replicon architecture for subsequent comparative analyses.

**Method S6. BGC prediction, clustering, diversity, and novelty analyse**

We analyzed 183 *Ktedonobacteria* genomes (Mount Zao MAGs/isolates plus public MAGs and cultured strains; Merged_NVS_bin65 excluded). To limit fragmentation bias, only scaffolds ≥5 kb were processed with antiSMASH v7.1.0[28] (bacterial mode, default), yielding 1,546 BGCs. BGCs were clustered with BiG-SLiCE v2.0[29] (cosine = 0.4) into 1,162 non-redundant gene cluster families (GCFs); GCF counts were overlaid on the phylogeny in iTOL[26]. This metric, recommended by Gavriilidou et al. (2022)[30], accounts for domain count bias and better reflects chemical similarity than raw Euclidean distances.

We classified a lineage as BGC-rich when its genome harbored ≥10 distinct GCFs. At the clade (family) level, we designated highly BGC-rich clades as those for which the mean per-genome GCF count across member genomes was ≥15 *(*optionally add: considering clades with ≥2 genomes).

These clade-level labels are independent of lineage-level labels. Each GCF was assigned to a biosynthetic class (NRP, polyketide, RiPP, terpene, saccharide, hybrid, others) based on antiSMASH/BiG-SLiCE annotations; for multi-BGC GCFs, the class of the longest BGC was used, and mixed classes were labeled "Hybrid." The same pipeline was applied to 33 Chloroflexota genomes, 115 *Streptomyces* type strains, and 1,135 Actinomycetota genomes (downloaded from NCBI GenBank with filters: type material, assembly level ≥CHR). GCF distributions were compared across lineages, genome types (MAG vs isolate), and clades using Wilcoxon rank-sum tests with Benjamini–Hochberg correction.

Novelty was assessed by three approaches: (i) BiG-SLiCE v1.1.1 against the BiG-FAM dataset (1,225,071 BGCs)[29, 31], considering distances >900 as novel[29]; (ii) BiG-SCAPE v1.1.2 against MiBIG v4.0[32], with d > 0.5 regarded as novel[33]; and (iii) sequence similarity to MiBIG entries based on antiSMASH KnownClusterBlast[28]. The same analyses were applied to 3,609 *Streptomyces* BGCs for comparison.

To evaluate diversity and redundancy, all BGCs from *Ktedonobacteria* and *Streptomyces* were clustered with BiG-SLiCE v2.0 at cosine thresholds 0.0/0.2/0.4/0.6, and the resulting GCF counts were compared across thresholds. Intra-genomic redundancy was assessed by comparing per-genome BGC counts to per-genome GCF counts at cosine 0.4 (clustering performed independently per genome).

For fine-scale patterns, each GCF carried both class and subclass annotations from BiG-SLiCE v2.0; multiple subclasses were retained (not merged) using the "Class_Subclass" format (e.g., "RiPP_bottromycin"). Subclass counts were Z-score normalized by column and visualized as a heatmap (pheatmap). Biosynthetic similarity among genomes was mapped by UMAP (R umap) with points annotated by taxonomy and habitat. Community structure was tested by PERMANOVA (adonis2, vegan; 999 permutations) using taxonomy (order, family) and ecosystem category as factors. Habitat associations of subclasses were modeled with GLMMs (binomial glmer, lme4) including lineage as a random intercept; odds ratios and Wald p-values were extracted (broom.mixed) and BH-corrected within habitat. Rare subclass–habitat pairs yielding non-estimable ORs were excluded from plots. All analyses/visualizations were run in R v4.2.0 with ggplot2, dplyr, umap, vegan, lme4, and broom.mixed.

**Method S7. Identification and comparative analysis of ECE-like contigs**

To identify candidate large ECEs distinct from the primary CHR, we defined them based on three criteria for initial screening.

(i) Only genomes sequenced with long-read technologies (e.g., PacBio). To further ensure high assembly quality, only genomes with ≤10 contigs were included in this analysis. (ii) Contigs ≥1 Mbp were selected as potential large replicons. This threshold was based on previous studies showing that chromids average 1.52 Mbp (median 1.26 Mbp), whereas megaplasmids average 0.77 Mbp (median 0.56 Mbp) (e.g., diCenzo & Finan, 2017[34]). (iii) Genome completeness was estimated using CheckM v1.0.12 with bacterial markers fixed to domain Bacteria[11]. Contigs were considered ECE-like contigs if their observed completeness was ≤0.2 of the expected value, defined as the contig length divided by total genome size. This operational threshold reflects the expectation that chromids lack most universal single-copy markers and therefore exhibit artificially low CheckM completeness relative to their size. By normalizing completeness to the expected value based on contig length, we aimed to distinguish large ECEs from fragmented genomes.

Terminal inverted repeats (TIRs) were detected by self-alignment with BLASTn v2.13.0. Strand asymmetry and replication structure were profiled from complete sequences with a custom Python script: Z-curve at 1-bp resolution (1,000-bp moving average) and normalized GC-skew in 1,000-bp windows. Replication-related genes (dnaA, dnaN, parA/repA, parB, repC) were identified with Pfam HMMs (HMMER), and origins were predicted with Ori-Finder 2022[35]. ParA/RepA proteins were aligned with MUSCLE v3.8.425[36], trimmed with trimAl v1.4[25] (-gt 0.5 -cons 60), and analyzed in IQ-TREE v2.3.6 (ModelFinder; Q.pfam+I+G4; 1,000 ultrafast bootstraps and 1,000 SH-aLRT).

Comparative genomics included pairwise ANIb with PyANI v0.3.0α (visualized as a clustered heatmap in R v4.3.1 with pheatmap); orthologous-gene presence/absence with Panaroo v1.5.1 (strict)[37], followed by Jaccard distances and PCoA (base R cmdscale); gene-set overlaps summarized with VennDiagram; and genome-wide distances with Mash v3.1[38] (k = 15; default sketch).

Proteins from CHR- and ECE-like contigs were annotated with eggNOG-mapper v2.1.12[39] (eggNOG v5.0.2; DIAMOND v2.1.9; --m diamond; tax_scope=bacteria). KEGG pathway presence/absence was plotted as a heatmap (pheatmap, R v4.3.1), and COG category proportions were compared between replicon types with ggplot2.

Pairwise comparisons between each CHR-like contig and its cognate ECE-like contig were performed with BLASTn v2.13.0 (megablast). Homologous regions were defined as ≥300 bp with ≥98% identity, and long terminal homology as ≥5 kb with ≥99% identity. Coordinates were extracted with custom Python scripts to generate Circos inputs. Features overlaid included BGCs (antiSMASH v7.1.0), rRNA genes (barrnap v0.9 via Prokka v1.14.6), and mobility-related genes parsed from eggNOG annotations ("integrase", "recombinase", "transposase"). Circos v0.69-9[40] was used for visualization.

To test whether mobility-associated genes are preferentially located inside or near biosynthetic gene clusters (BGCs), we quantified overlap and proximity between antiSMASH-predicted BGC intervals and mobility-gene intervals (integrase, recombinase, and transposase annotations parsed from eggNOG). Coordinates were handled as half-open intervals $[start, end)$ on each replicon.

For each replicon (CHR-like or ECE-like) in the 20 long-read genomes analyzed (Fig. 5), we defined: (i) the BGC-covered length $W$ (bp; union of BGC intervals), (ii) the flanking length $F$ (bp; union of the 2×$L$-bp windows immediately outside each BGC boundary, excluding any bases within BGCs and merging overlaps among flanks), and (iii) the total replicon length $G$ (bp). Each mobility interval was classified as within (overlapping any BGC base), flank (not within, but overlapping the flank region), or outside (neither within nor flank). Length-corrected enrichment was evaluated with a binomial model in which the null probability of a mobility interval falling in a region equals the region length divided by replicon length ($p_{\text{within}} = W/G;\ p_{\text{flank}} = F/G$). For each window size $L$ (2, 5, 10, and 20 kb), we summed mobility intervals across replicons ($N$) and counted those within ($k_{\text{within}}$) and in flank ($k_{\text{flank}}$). One-sided binomial tests were used to assess depletion within BGCs $\Pr[X \leq k_{\text{within}}]$ and enrichment in flanks $\Pr[X \geq k_{\text{flank}}]$. Effect sizes were reported as fold enrichment $FE = (k/N)/(region\_length/G)$, where $FE > 1$ indicates enrichment and $FE < 1$ indicates depletion.

As a robustness check that does not rely on the binomial assumptions, we performed a within-replicon permutation test: mobility intervals were randomly relocated on the same replicon while preserving each interval length and keeping all intervals within replicon boundaries; this was repeated 1,000 times per $L$ to generate null distributions of $k_{\text{within}}$ and $k_{\text{flank}}$. Permutation $p$-values were computed as $(r+1)/(n_{\text{perm}}+1)$, where $r$ is the number of permutations with counts at least as extreme as observed (≥ for flank enrichment; ≤ for within depletion).

To evaluate whether mobility loci outside BGCs are biased toward BGC proximity, we calculated, for each mobility interval classified as outside, the minimum distance (bp) to the nearest BGC boundary on the same replicon. Distances were summarized into bins (0–2, 2–5, 5–10, 10–20, and ≥20 kb) and as cumulative thresholds (<2, <5, <10, and <20 kb). Observed counts were compared with the same permutation scheme to obtain enrichment ratios (observed / null mean) and one-sided permutation $p$-values for each bin and threshold.

For BGC subclass distributions, BiG-SLiCE v2.0 subclass labels (e.g., NRP_siderophore, Polyketide_T1PKS) were retained without aggregation. Per-replicon presence/absence and counts were compiled in R v4.3.1 (tidyverse v2.0.0; vegan v2.6-4). Bray–Curtis dissimilarities were used for PCoA (ape v5.7) to compare subclass composition, with separation by replicon type (CHR-like contig vs ECE-like contig) tested by PERMANOVA (adonis2, 999 permutations). For paired CHR-like contig/ECE-like contig comparisons, subclasses present in ≥3 genomes in either replicon type were tested with exact McNemar's tests and Benjamini–Hochberg correction; results were shown as barplots and heatmaps (pheatmap v1.0.12).

https://doi.org/10.1093/bioinformatics/btv638

24. Yabe S et al. Dictyobacter aurantiacus gen. nov., sp. nov., a member of the family Ktedonobacteraceae,

isolated from soil, and emended description of the genus Thermosporothrix. *Int J Syst Evol Microbiol* 2017;**67**:2615–2621. https://doi.org/10.1099/ijsem.0.001985

32. Zdouc MM et al. MIBiG 4.0: advancing biosynthetic gene cluster curation through global collaboration.

*Nucleic Acids Res* 2025;**53**:D678–D690. https://doi.org/10.1093/nar/gkae1115
